# Point mutation in a virus-like capsid drives symmetry reduction to form tetrahedral cages

**DOI:** 10.1101/2024.02.05.579038

**Authors:** Taylor N. Szyszka, Michael P. Andreas, Felicia Lie, Lohra M. Miller, Lachlan S. R. Adamson, Farzad Fatehi, Reidun Twarock, Benjamin E. Draper, Martin F. Jarrold, Tobias W. Giessen, Yu Heng Lau

## Abstract

Protein capsids are a widespread form of compartmentalisation in nature. Icosahedral symmetry is ubiquitous in capsids derived from spherical viruses, as this geometry maximises the internal volume that can be enclosed within. Despite the strong preference for icosahedral symmetry, we show that simple point mutations in a virus-like capsid can drive the assembly of novel symmetry-reduced structures. Starting with the encapsulin from *Myxococcus xanthus*, a 180-mer bacterial capsid that adopts the well-studied viral HK97 fold, we use mass photometry and native charge detection mass spectrometry to identify a triple histidine point mutant that forms smaller dimorphic assemblies. Using cryo-EM, we determine the structures of a precedented 60-mer icosahedral assembly and an unprecedented 36-mer tetrahedron that features significant geometric rearrangements around a novel interaction surface between capsid protomers. We subsequently find that the tetrahedral assembly can be generated by triple point mutation to various amino acids, and that even a single histidine point mutation is sufficient to form tetrahedra. These findings represent the first example of tetrahedral geometry across all characterised encapsulins, HK97-like capsids, or indeed any virus-derived capsids reported in the Protein Data Bank, revealing the surprising plasticity of capsid self-assembly that can be accessed through minimal changes in protein sequence.

**Significance statement:** Viral capsids are cage-like protein assemblies that preferentially adopt icosahedral symmetry to maximise their internal volume for housing genetic material. This icosahedral preference extends to encapsulins, a widespread family of bacterial protein cages which evolved from viral capsids. Counter to this fundamental geometric preference, the formation of well-defined tetrahedral cages from a single amino acid substitution in an encapsulin reveals the surprising geometric flexibility of a common viral protein fold. These findings suggest that protein oligomerisation is far more permissive than intuitively expected, where serendipitous interactions between proteins arising from minimal mutations can cascade to form vast architectural changes. The ability to redesign protein architectures through simple mutations should enable biotechnological advances in vaccine development, drug delivery, and enzymatic biomanufacturing.

## Introduction

Cage-like protein assemblies are a ubiquitous structural design feature in biology. Viral capsids are the most well-studied example, serving as a protective shell that houses the viral genome.^(1)^ Prokaryotes use protein cages as organelles, hosting enzymes within their interior to confine metabolic reactions into organised intracellular spaces.^(2–6)^ In eukaryotes, Arc proteins form virus-like capsids that transport RNA between neurons,^(7, 8)^ while in all organisms, ferritin protein cages serve as essential iron storage compartments.^(9)^ Beyond naturally occurring capsids, *de novo* designed cages have been used as scaffolds for antigen display to create new vaccines.^(10, 11)^

A common feature across all cage-like assemblies is their high degree of structural symmetry, driven by the principle of maximising genetic economy.^(12)^ In the simplest case, one gene is sufficient to encode the formation of an entire capsid, where multiple copies of the encoded protomer self-assemble into closed hollow structures with spherical morphology. Depending on the rotational axes of symmetry present, capsids with overall spherical morphology can be classed as having symmetry that is icosahedral (5/3/2 point symmetry, 60 asymmetric units), octahedral (4/3/2 point symmetry, 24 asymmetric units), or tetrahedral (3/2 point symmetry, 12 asymmetric units), listed in decreasing order of symmetry. Capsids with icosahedral symmetry are particularly prevalent in nature, as this geometric arrangement maximises the volume enclosed by a capsid from a fixed amount of genetic coding information.^(12)^ These geometric preferences are accentuated in viral capsids that are not enveloped by a lipid membrane, as the capsid must maintain a robust morphology to function as the physical outer layer.^(13)^

Even when structural changes occur upon introducing mutations to a capsid protein, spherical morphology and symmetry group are typically preserved. The widespread family of capsids with protomers that adopt the HK97 viral fold provides an archetypal example of this phenomenon. Originally named after the HK97 bacteriophage capsid, the HK97 fold is found in all Herpesviruses, tailed bacteriophages, some archaeal viruses, and prokaryotic encapsulin nanocompartments, forming different icosahedral capsids that range from 18 nm to 180 nm in diameter.^(14, 15)^ The size is largely dependent on the number of protomers involved in the assembly, which can be defined by the triangulation number (*T*, where 60×*T* is the number of protomers).^(16, 17)^ Icosahedral capsids typically contain pentameric facets that provide the necessary curvature to form a closed polyhedron, along with optional hexamers that sit between pentameric vertices to extend the size of the overall capsids. In most mutational studies on icosahedral capsids, the introduced mutations compromise hexamer formation and lead to small pentamer-only *T*=1 capsids that still have spherical morphology and icosahedral symmetry, whereas mutations that compromise pentamer formation can sometimes lead to elongated filamentous assemblies due to insufficient curvature.^(13)^ Beyond these two ordered mutant assemblies and other irregular mis-assemblies, no other examples of defined structural polymorphism have been reported over many decades of structural studies on the HK97-like family of capsids.^(14, 15)^ More generally, there are very few known examples of structural polymorphism in virus-like capsids that feature a shift to significantly different yet well-defined morphologies or symmetries.^(13)^

Herein, we report the serendipitous discovery of simple point mutants of an HK97-fold encapsulin protein that forms assemblies with unprecedented tetrahedral symmetry and morphology. Encapsulins are a widespread family of prokaryotic protein cages that house specific enzymes as cargo within their interior, each with diverse native functions such as iron storage or response to oxidative stress.^(2, 3, 18–20)^ Cargo encapsulation is mediated by a targeting peptide sequence at the C-terminus of each cargo protein, where the peptide binds a pocket on the interior surface of the encapsulin protomer.^(19, 21–24)^ Encapsulins are exclusively icosahedral, as with all HK97-like capsids.^(14, 15)^ Neither native nor mutant cages with the HK97 fold have ever been reported to adopt lower symmetry octahedral or tetrahedral forms in the Protein Data Bank.

## Results

### Pore-lining point mutations generate smaller dimorphic assemblies

Our investigation began by constructing pore mutants of the encapsulin protein cage from *Myxococcus xanthus* (MxEnc), which natively forms spherical 180-mer assemblies with *T*=3 icosahedral symmetry. When recombinantly expressed in the absence of any packaged cargo, MxEnc also forms a minor proportion of smaller 60-mer assemblies with *T*=1 icosahedral symmetry.^(20, 22)^ Given that we had previously observed extraordinary robustness of cage assembly across a diverse range of pore mutants of the encapsulin from *Thermotoga maritima*,^(25)^ we hypothesised that pore mutation would have minimal impact on MxEnc assembly, while enhancing the permeability of the cage to substrates and products of encapsulated enzymatic reactions in a bioengineering context.^(26–28)^

The first cage we constructed was the triple point mutant K199H/T200H/G201H, henceforth referred to as 3×His-MxEnc. These three residues are positioned at the centre of the apical A-domain loop of MxEnc that lines the exterior opening of the pentameric and hexameric pores (Figure 1a). Histidine was chosen due to its moderately-sized imidazole side chain with the potential for achieving charge-based gating of diffusion through the pore with dependence on the protonation state.

**Figure 1.**
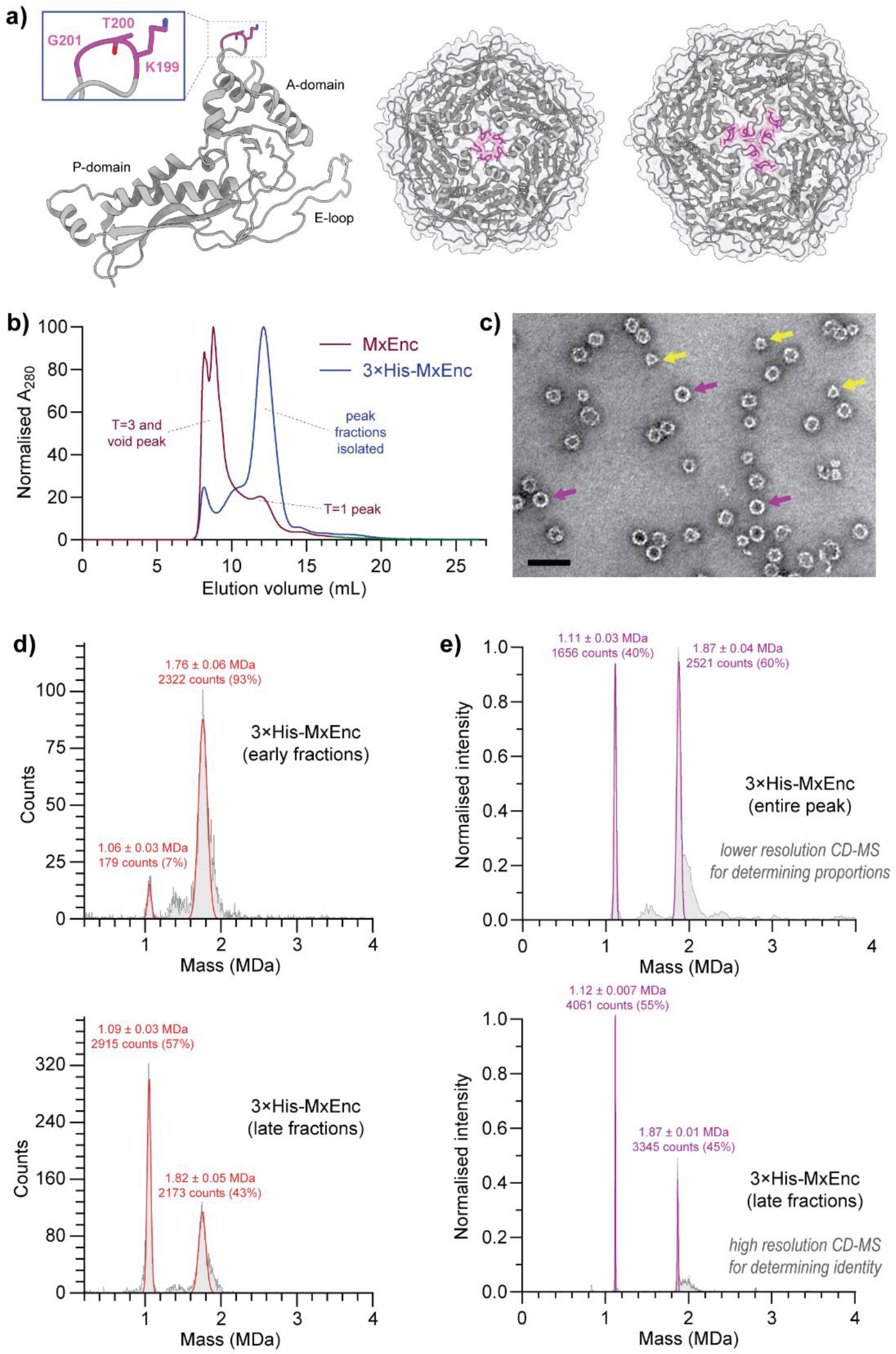
Point mutation of the *M. xanthus* encapsulin protein cage (MxEnc) results in smaller dimorphic assemblies. **a)** Published structure of wild-type MxEnc (PDB 7S20),^(22)^ highlighting the three residues of the A-domain loop that are mutated to histidine in the encapsulin mutant 3×His-MxEnc. These residues line the pore of the pentamers and hexamers in the wild-type assembly. An individual protomer (left), pentamer (middle), and hexamer (right) are shown with the three mutated residues highlighted in magenta. **b)** Size-exclusion chromatography on a Superose 6 Increase column indicated a clear shift towards smaller assemblies. **c)** Negative stain transmission electron microscopy showed the presence of two assemblies – a smaller assembly with clear triangular features visible at some angles (yellow arrows), and a larger assembly with a more spherical morphology (purple arrows). Scale bar = 50 nm. **d)** Mass photometry indicated the presence of two distinct assemblies that were differentially enriched in different fractions from size-exclusion chromatography. **e)** Native CD-MS confirmed the existence of a 36-mer and a 60-mer as the two dominant species isolated. Errors are reported as the standard deviation.

During the purification of recombinantly produced 3×His-MxEnc, we observed an unexpected significant shift towards smaller assemblies in size-exclusion chromatography when compared to wild-type MxEnc (Figure 1b). Unlike the expected profile of a major peak corresponding to the *T*=3 assembly and a minor peak for the *T*=1 assembly, the major size-exclusion peak for 3×His-MxEnc eluted at a volume range expected for a smaller *T*=1 assembly. Conducting transmission electron microscopy (TEM) on negatively stained samples of 3×His-MxEnc, we observed spherical particles with a diameter of 24 nm as expected for a *T*=1 symmetric assembly, alongside smaller non-spherical particles with well-defined triangular features corresponding to a smaller unknown assembly (Figure 1c).

After size-exclusion chromatography, we analysed fractions containing pure 3×His-MxEnc by mass photometry, revealing the co-existence of two well defined and kinetically stable assemblies that were differentially enriched across different size-exclusion fractions (Figure 1d). At ∼1.1 MDa and ∼1.8 MDa respectively, both assemblies were far smaller than the expected mass of 5.7 MDa for the wild-type *T*=3 MxEnc. To identify the oligomeric state of the assemblies and quantify their relative proportions in the overall sample, we subsequently used the greater resolving power of native charge detection mass spectrometry (CD-MS), an emerging technique where simultaneous measurements of the charge and the mass-to-charge ratio provide single particle mass information for large polymorphic assemblies (Figure 1e).^(29, 30)^ Using a pooled sample of all encapsulin-containing fractions, CD-MS measurements indicated that the larger assembly had a mass of 1.87 ± 0.01 MDa, consistent with a *T*=1 icosahedral 60-mer assembly. At two-thirds the abundance of the larger assembly based on particle counts in CD-MS, the smaller assembly with a mass of 1.12 ± 0.007 MDa was consistent with a novel 36-mer assembly. There was no evidence of any *T*=3 assemblies reminiscent of wild-type MxEnc by TEM, mass photometry, or native CD-MS.

### Cryo-EM reveals the structure of a novel tetrahedral protein cage

We elucidated the structure of 3×His-MxEnc in its two co-existing assembly states by single particle cryo-electron microscopy (cryo-EM) analysis, confirming their respective identities as a 60-mer assembly with *T*=1 icosahedral symmetry and a 36-mer assembly with tetrahedral symmetry and morphology (Figure 2). The structure of the *T*=1 icosahedral assembly broadly resembles the published *T*=1 structure that arises exclusively from recombinant MxEnc expression in the absence of co-expressed cargo (Figure 2a).^(20, 22)^ Meanwhile, the tetrahedral assembly is unprecedented across all encapsulins (Figure 2b), or indeed any capsids with the HK97-fold, or indeed any known protein cages with a virus-like fold that have been reported in the Protein Data Bank to date (Supplementary File 1).

**Figure 2.**
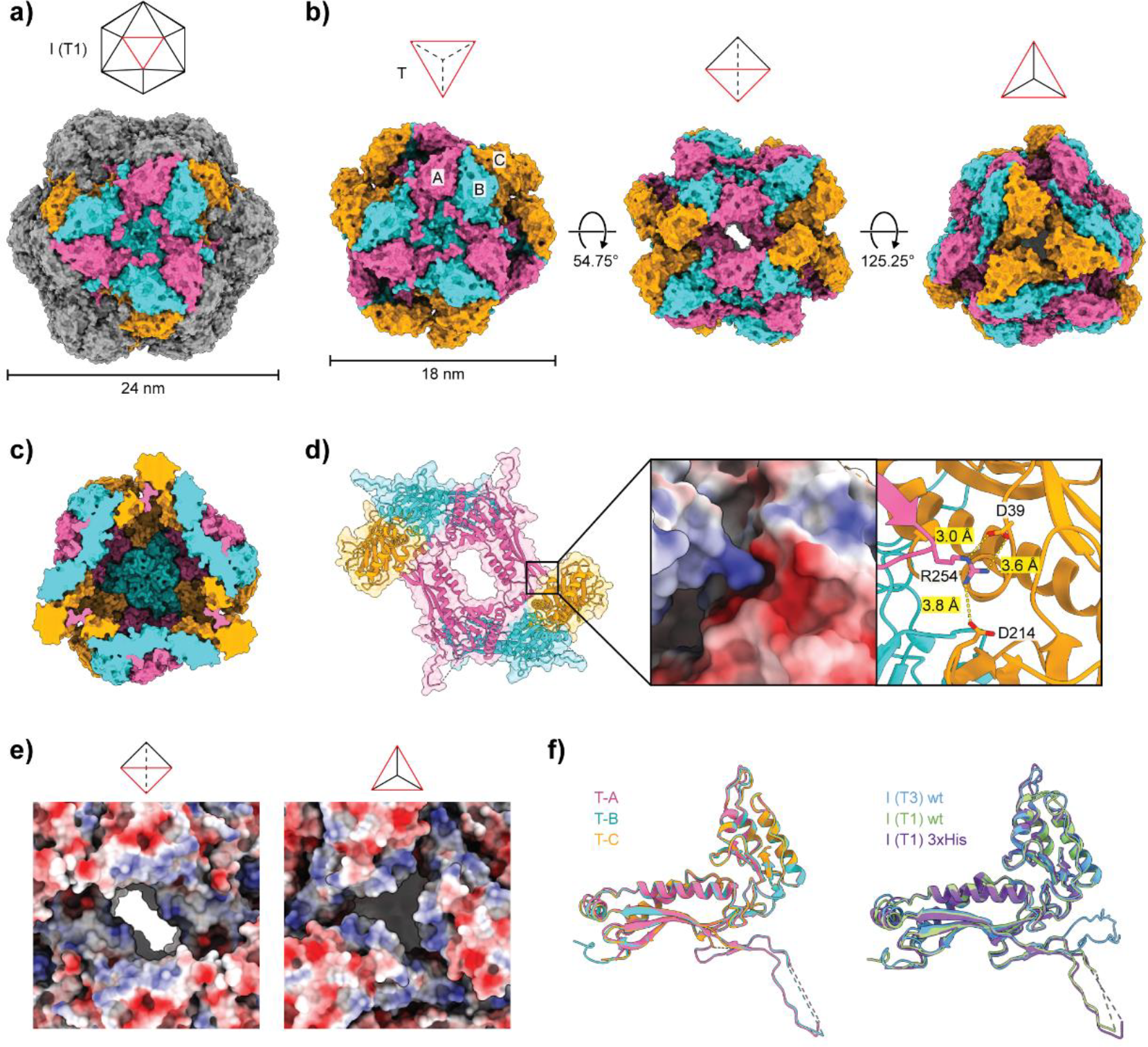
Cryo-EM structures of the dimorphic assembly states of 3×His-MxEnc. **a)** The *T*=1 icosahedral form of 3×His-MxEnc closely resembles the *T*=1 structure of recombinant wild-type MxEnc produced in the absence of cargo.^(22)^ **b)** The tetrahedral form of 3×His-MxEnc is comprised of three non-equivalent protomers A-C in the asymmetric unit. The three-fold symmetric interaction at the centre of each triangular face is conserved, while there are novel two-fold and three-fold interactions at the edges and vertices, respectively. **c)** The interior volume of the tetrahedron is reduced 6.5-fold in comparison to the *T*=1 icosahedral assembly and 40-fold compared to the wild-type *T*=3 cage. **d)** A novel two-fold symmetric interaction between two offset trimers drives tetrahedron assembly. R254 from protomer A forms an electrostatic interaction with D39 and D214 from protomer C, thus occupying the site where the targeting peptide of encapsulated cargo usually binds. **e)** The novel two-fold interaction creates a large and elongated pore at each edge, while the novel three-fold interaction at the vertices is highly flexible and is likely less crucial to assembly. **f)** In the tetrahedron, the E-loop of protomer C is completely unresolved and likely flexible. The E-loops of protomers A and B adopt the same orientation as found in the *T*=1 icosahedral forms of both 3×His-MxEnc and wild-type MxEnc.

The tetrahedral cage is assembled from four largely flat triangular faces, each comprised of three copies of the asymmetric unit that contains three distinct protomers – A, B, and C (Figure 2b). The three-fold symmetric interaction at the centre of each triangular face is the only interaction that is conserved from the wild-type MxEnc assembly. There is an anti-clockwise rotational offset of ca. 13° to each triangular face, such that the overall morphology deviates slightly from an ideal tetrahedron, with new two-fold and three-fold symmetric interactions at the edges and vertices, respectively. As a result of symmetry reduction, the interior volume of the tetrahedral cage of 300,040 Å^3^ is significantly less than that of the *T*=1 icosahedral cage (1,944,000 Å^3^) or the wild-type *T*=3 cage (11,980,000 Å^3^) (Figure 2c).

Tetrahedral cage assembly is driven by a novel two-fold symmetric interaction which occurs at the targeting peptide site where enzymatic cargo is normally bound in the native encapsulin (Figure 2d).^(22, 23)^ Along each two-fold symmetric edge of the tetrahedron, a pair of trimers meet in a staggered alignment to form an offset hexameric arrangement, with the histidine mutations at the A-domain loop likely disfavouring the formation of standard hexameric or pentameric interactions. Instead, a novel key electrostatic interaction is formed between the side-chain of R254 from the P-domain of protomer A, and D39 and D214 on the interior surface of a neighbouring protomer C. In wild-type MxEnc, these same two aspartate residues mediate cargo encapsulation by interacting with a key arginine residue from the *C*-terminal targeting peptide region of the native enzymatic cargo.^(22, 23)^ The result of this novel two-fold interaction is a more porous structure relative to the wild-type MxEnc, with an elongated two-fold symmetric pore with approximate dimensions of 15×30 Å at each edge of the tetrahedron (Figure 2e).

The novel three-fold symmetric interaction at the vertices is likely less important for tetrahedron assembly (Figure 2e). There is significant flexibility at the vertices as seen by the drop in local resolution around this region in the tetrahedral cryo-EM density (SI Figure S5.3). Furthermore, the entire E-loop (residues 45-82) of each protomer C that flanks the vertices could not be resolved and is likely to be highly flexible (Figure 2f). Meanwhile, the E-loops of protomers A and B adopt a similar orientation to that observed in both the *T*=1 wild-type MxEnc assembly and the *T*=1 assembly state of 3×His-MxEnc.

### Filled T=1 icosahedral assemblies are generated upon cargo expression

Co-expression of 3×His-MxEnc and the fluorescent protein mNeonGreen with a *C*-terminal targeting peptide as cargo (mNeonTP) resulted in the formation of a filled *T*=1 icosahedral assembly and loss of the tetrahedral assembly (Figure 3). During size-exclusion chromatography, we observed a peak that eluted at the expected size range for *T*=1 assemblies, along with a higher order aggregate peak that eluted in the unretained peak of the Superose 6 Increase column (Figure 3a). SDS-PAGE analysis of size-exclusion fractions indicated that both peaks contained encapsulin and fluorescent cargo (SI Figure S2.5). Native PAGE analysis showed that the *T*=1 icosahedron was the dominant assembled species, while no band for the tetrahedral assembly was observed (Figure 3b). In-gel fluorescence was observed for the *T*=1 band, confirming the presence of the mNeonTP cargo with an average loading density of four cargo molecules per encapsulin assembly as determined by relative UV-Vis absorbance.

**Figure 3.**
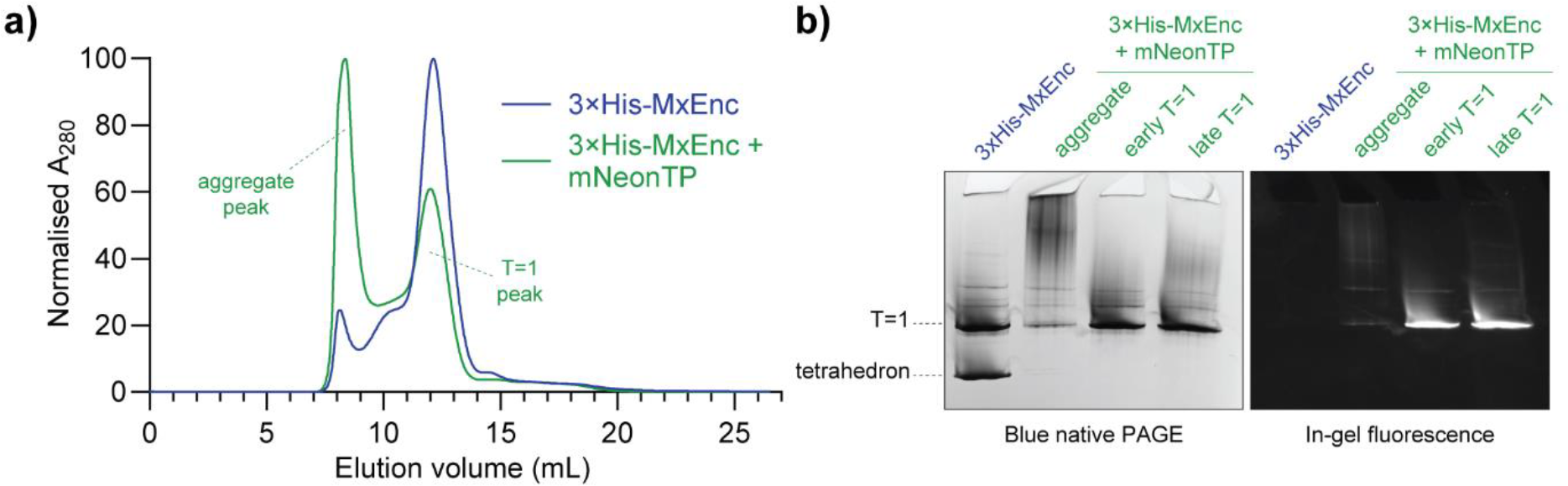
The *T*=1 icosahedral assembly is the dominant species in the presence of cargo. **a)** Size-exclusion chromatography on a Superose 6 Increase column indicates that co-expression of 3×His-MxEnc with mNeonTP cargo causes a slight leftwards shift in the *T*=1 peak, as well as the formation of large aggregates that elute in the unretained peak. **b)** Native PAGE analysis shows a dominant *T*=1 assembly with accompanying in-gel fluorescence when mNeonTP cargo is present, while there is no evidence of tetrahedral assemblies.

As the wild-type MxEnc exclusively forms *T*=3 icosahedral assemblies when co-expressed with cargo,^(20, 22, 23)^ the unexpected observation of filled *T*=1 assemblies for 3×His-MxEnc provides further evidence to suggest that the histidine mutations at the A-domain loop destabilise the ability to form hexameric facets, which are required for *T*=3 assemblies but not present in *T*=1 assemblies. Meanwhile, the absence of any filled tetrahedral assemblies is caused by competition from the targeting peptide of the mNeonTP cargo, occupying the D39/D214 interaction site that mediates the novel two-fold interaction in the tetrahedron, providing further confirmation of the importance of the electrostatic interaction with R254.

### Tetrahedral cage assembly is not exclusive to histidine mutants

To assess the impact of point mutations beyond histidine, alternative triple point mutants of residues 199-201 to alanine, lysine and aspartic acid were generated, forming 3×Ala-MxEnc, 3×Lys-MxEnc, and 3×Asp-MxEnc, respectively. These constructs were purified using the same workflow as the original histidine mutant 3×His-MxEnc.

The triple alanine mutant 3×Ala-MxEnc showed similar assembly behaviour to the original histidine mutant 3×His-MxEnc. During purification, both samples had similar size-exclusion chromatography traces (Figure 4a and 4b), with a major peak at the retention time corresponding to tetrahedral and *T*=1 icosahedral assemblies. Native PAGE analysis of fractions from size-exclusion chromatography confirmed the presence of both assembled states (Figure 4c), with matching data obtained from mass photometry measurements (Figure 4d) and negative stain TEM (Figure 4e).

**Figure 4.**
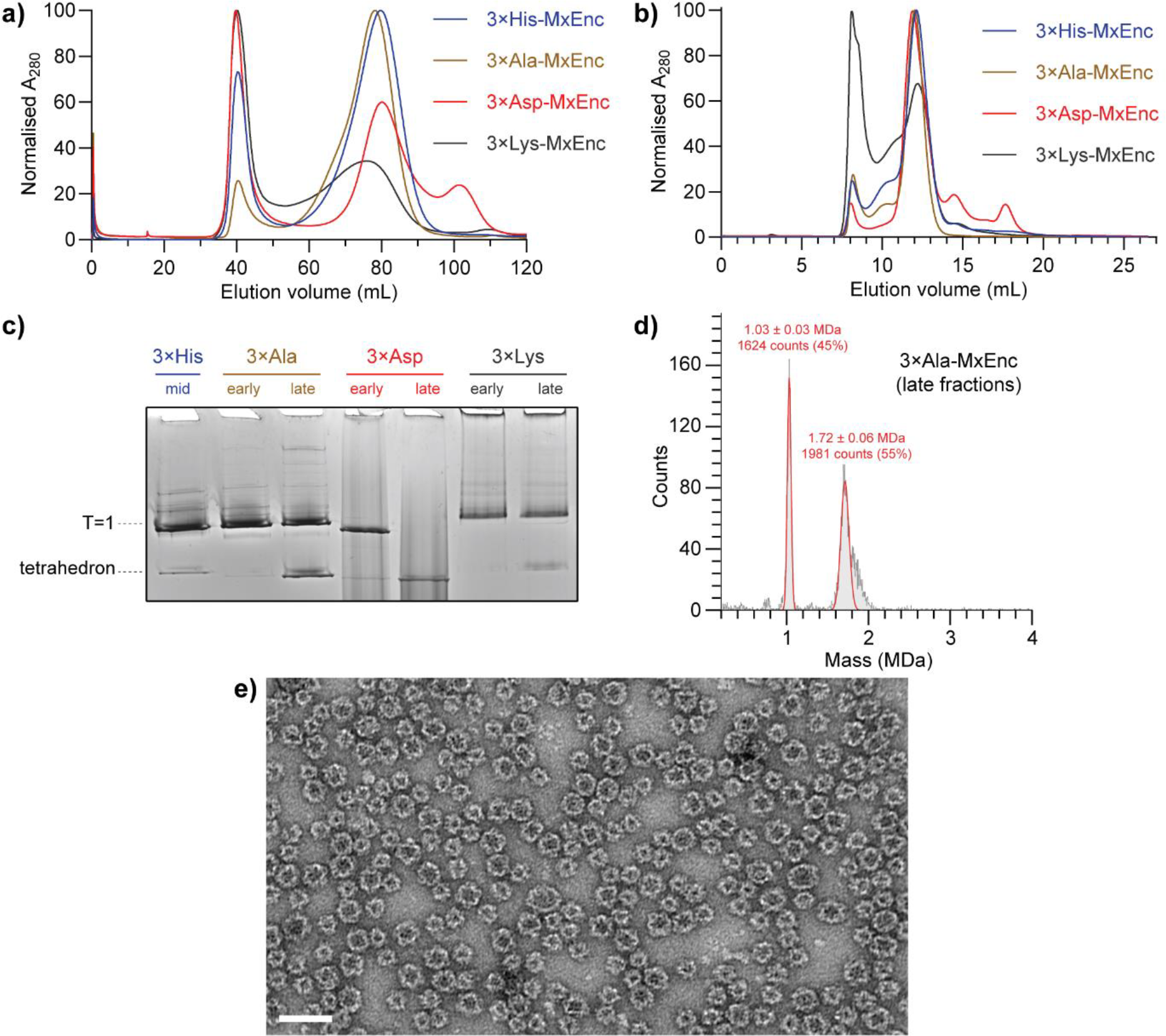
Tetrahedral cages can be formed by point mutation to various amino acids. **a)** Initial low-resolution size-exclusion chromatography on a Sephacryl S-500 column shows that the 3×His and the 3×Ala mutants have similar profiles, while the charged 3×Asp and 3×Lys mutants are more polydisperse. Fractions in the ∼80 mL peak were isolated for downstream purification. **b)** Subsequent higher-resolution size-exclusion chromatography on a Superose 6 Increase column revealed similar trends, where the 3×His and 3×Ala mutants were similar, while the 3×Asp and 3×Lys mutants were more polydisperse. **c)** Blue native PAGE analysis of early or late fractions from the main peak eluted from the Superose 6 Increase column for 3×Ala, 3×Asp and 3×Lys mutants, along with a middle fraction of the original 3×His mutant as a size standard. Tetrahedral and *T*=1 icosahedral bands can clearly be observed for 3×Ala-MxEnc. There is evidence of a tetrahedral band for 3×Asp-MxEnc although smearing for suggests polydispersity and instability. A faint tetrahedral band can be observed for 3×Lys-MxEnc. **d)** Mass photometry confirms that the 3×Ala mutant can form tetrahedral and *T*=1 icosahedral assemblies. **e)** Negative stain transmission electron micrograph of 3×Ala-MxEnc provides visual evidence of tetrahedral and *T*=1 icosahedral assemblies. Scale bar = 50 nm.

Mutation to charged residues in 3×Lys-MxEnc and 3×Asp-MxEnc led to an increase in aberrant assembly behaviour. The positively charged mutant 3×Lys-MxEnc showed a significant increase in larger polydisperse protein assemblies that eluted in the unretained peak during size-exclusion chromatography on a Superose 6 Increase column (Figure 4a and 4b). Nevertheless, there was evidence of *T*=1 and a minor proportion of tetrahedral assemblies by Native PAGE analysis (Figure 4c). The negatively charged mutant 3×Asp-MxEnc appeared to be highly polydisperse during initial low-resolution size-exclusion chromatography steps on a Sephacryl S-500 column (Figure 4a). When the species in the *T*=1 size range were further separated on a Superose 6 Increase column, native PAGE analysis showed prominent streaking of the sample that indicated general instability of the mutant, although bands were still visible for *T*=1 and tetrahedral assemblies (Figure 4c). Polydispersity was also observed by mass photometry (SI Figure S4.1b). The aberrant nature of these assemblies is likely due to electrostatic repulsion from the clustering of the triple mutations at the A-domain loops around axes of symmetry (tetrahedral two-fold, icosahedral five-fold).

### Single point mutation is sufficient to generate tetrahedral cages

Formation of the tetrahedral assembly was observed in three independent single point mutants at residues 199-201 (K199H-MxEnc, T200H-MxEnc, G201H-MxEnc). During initial size-exclusion chromatography on a Sephacryl S-500 column (Figure 5a), there was a clear progressive shift towards smaller assemblies when going from K199H to G201H, where K199H-MxEnc had the most similar profile to wild-type MxEnc, and G201H-MxEnc had the most similar profile to the original triple mutant

**Figure 5.**
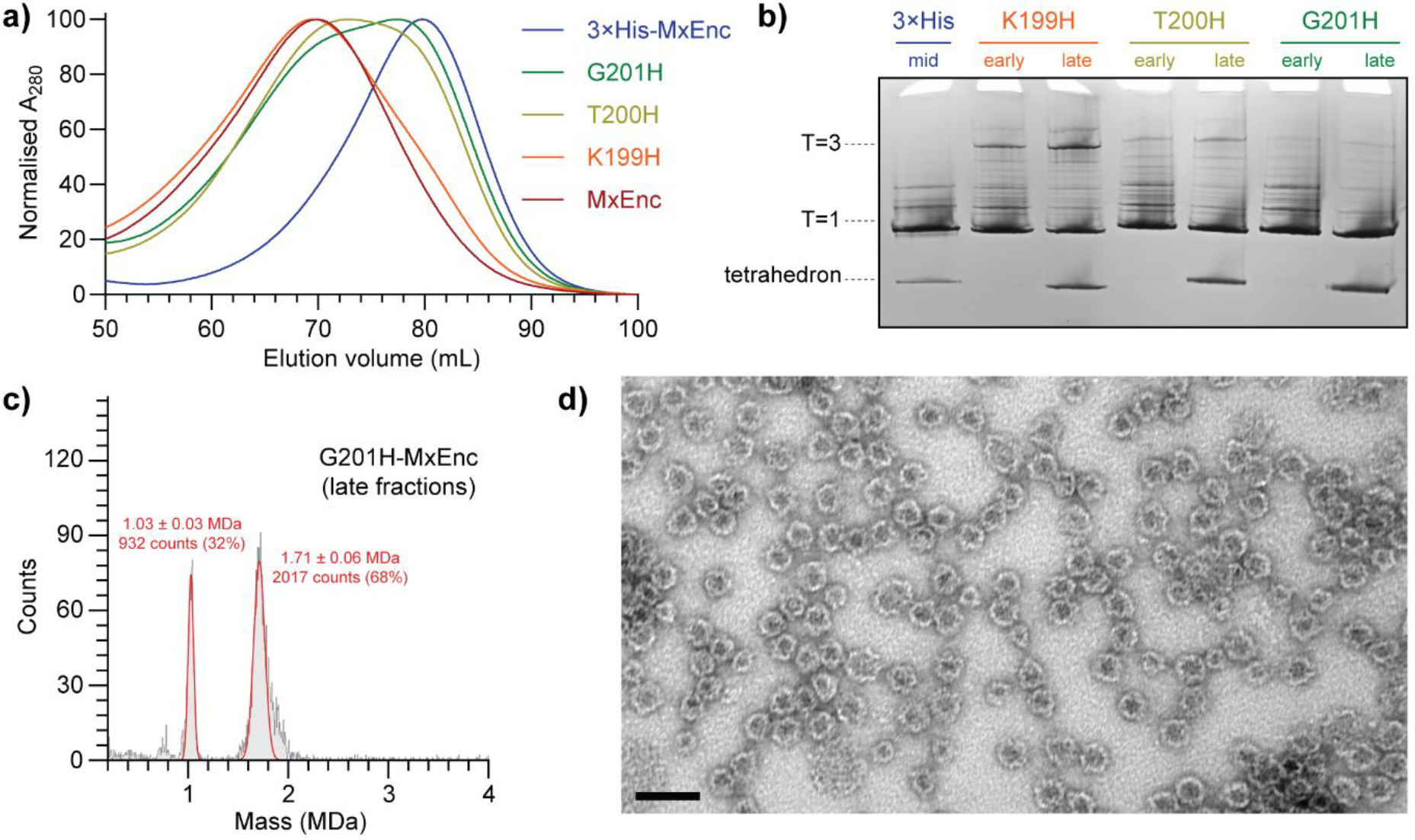
Tetrahedral cages can be generated by a single point mutation. **a)** Progressive single point mutations to histidine for residues 199-201 show a clear progressive shift towards the smaller assembly state by size-exclusion chromatography on a Sephacryl S-500 column, with the G201H mutant closest in behaviour to the original 3×His mutant. **b)** A tetrahedral band can be observed in blue native PAGE analysis of all single point mutants, along with the progressive shift towards smaller assemblies going from K199H to G201H. **c)** Mass photometry confirms that the G201H mutant can form tetrahedral and *T*=1 icosahedral assemblies. d) Negative stain transmission electron micrograph of G201H-MxEnc provides visual evidence of tetrahedral and *T*=1 icosahedral assemblies. Scale bar = 50 nm.

3×His-MxEnc. Analysing early and late fractions from the main peak by blue native PAGE (Figure 5b), along with a sample of 3×His-MxEnc as a size standard, there was clear evidence of both the tetrahedral and *T*=1 icosahedral assembly in later fractions of all three mutants. Consistent with the size-exclusion chromatography data, native PAGE of G201H was devoid of any distinct strong bands for higher-order assemblies, while K199H showed evidence of the larger *T*=3 assembly state, and the wild-type MxEnc only showed evidence of the known *T*=3 and *T*=1 icosahedral assemblies (SI Figure S3.1). Mass photometry measurements confirmed the presence of both species in all single point mutants (Figure 5c and SI Figure S4.1). Negative stain TEM of G201H provided visual confirmation of both assemblies (Figure 5d).

Taken together, tetrahedral cage formation appears to be most favoured by mutation at G201. Indeed, G201H is the least conservative mutation, causing major changes in backbone flexibility, sterics, and charge. These factors should result in a more rigid A-domain loop while introducing greater steric hindrance and electrostatic repulsion, thus disfavouring hexamer and pentamer formation and resulting in symmetry reduction to the tetrahedral cage. In addition, evolutionary conservation analysis using ConSurf indicates that G201 is the most highly conserved of the three mutated pore residues, followed by T200, with K199 being least conserved (SI Figure S7.1).^(31)^ While there are many factors that could contribute to these trends, the importance of the G201 position is nevertheless reinforced by this conservation analysis.

### Surface tiling diagrams classify the geometry of polymorphic encapsulin assemblies

The surface tilings of the different cage architectures can be rationalised as different embeddings of polyhedral surfaces into the same planar grid, given here by a gyrated hexagonal lattice (Figure 6).^(32)^ The wild-type *T*=3 icosahedral MxEnc assembly corresponds to an embedding of an icosahedral surface (Figure 6a). With the formation of hexamers disfavoured by the apical A-domain loop mutations to histidine, the resulting pentamer-only icosahedral cage can be represented as an embedding of a smaller, rescaled icosahedral net (Figure 6b), that now represents a *T*=1 surface architecture. It is also possible to embed tetrahedral and octahedral surfaces into the gyrated hexagonal lattice in different ways.^(32)^ Some of these options would require the formation of trimeric and tetrameric clusters with rotational symmetry around the A-domain loop (SI Figure S8.1), which have not been observed for this protomer, perhaps as the angle of the required inter-protomer interactions would be very narrow, making these structures unlikely to occur in our context. By contrast, the tetrahedral embedding with offset hexamers is an experimentally observed protomeric arrangement (Figure 6c).

**Figure 6.**
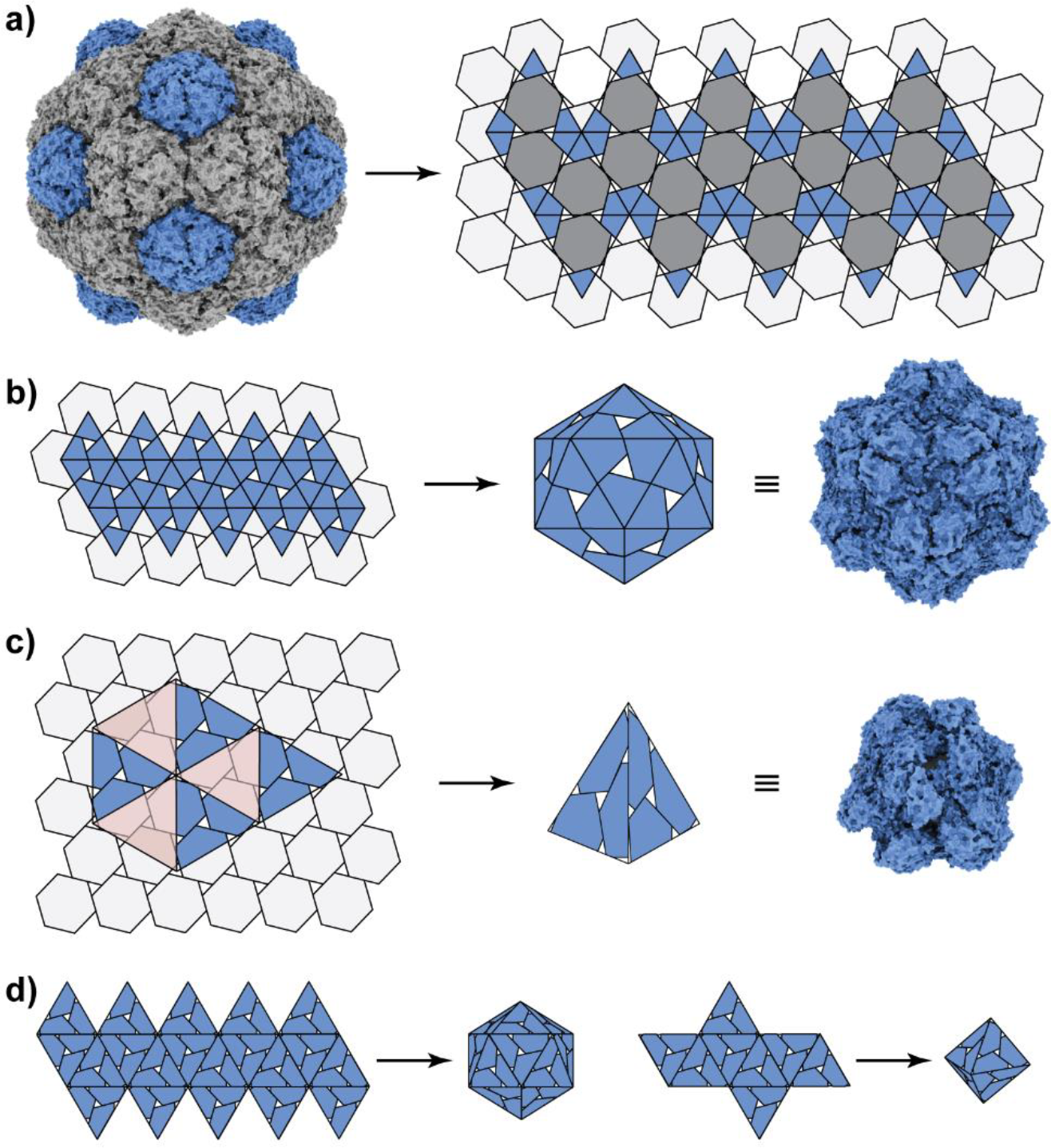
Surface tilings of encapsulin cages with icosahedral and tetrahedral symmetry. **a)** The tiling associated with the surface architecture of wild-type MxEnc is a gyrated hexagonal lattice. **b)** A planar embedding of a smaller icosahedral net into the gyrated hexagonal lattice generates the surface structure of the *T*=1 cage. **c)** A planar embedding of a tetrahedral net into the gyrated hexagonal lattice, aligning three-fold symmetry axes and removing the translucent red triangles, generates the surface structure of the observed tetrahedral encapsulin cage. **d)** Triangular faces of the tetrahedral cage may potentially be able to form novel particles with icosahedral and octahedral symmetry.

The tetrahedral embedding can be constructed geometrically by aligning the three-fold symmetry axes of a tetrahedron with those of the gyrated hexagonal lattice, i.e. placing the tetrahedral vertices where three hexagons meet (Figure 6c). To create the required three-fold symmetry axis, triangles shown in translucent red colour must be removed from the lattice when creating the 3D model from the planar lattice. This new embedding generates a cage with tetrahedral symmetry that captures the structural characteristics of the experimentally observed mutant tetrahedral cage. Based on the triangular faces of this tetrahedral embedding, it is theoretically possible to create alternative nets with octahedral or icosahedral symmetry (Figure 6d). In analogy to the tetrahedral particle, these cage structures would be formed entirely from offset hexamers. Instead of meeting in groups of three at the vertices of the tetrahedron, offset hexamers would meet in groups of four or five around the E-loop in these hypothetical new cages. Given the significant flexibility experimentally observed at the vertices of the mutant tetrahedron, it may be feasible to redesign the beta-turn loop that contains R254 and mediates the key two-fold symmetric interaction at the offset hexamers, potentially giving rise to new assembled geometries.

## Discussion

The mutant encapsulins generated in this study represent the first report of tetrahedral assemblies that have been constructed from protomers with a viral fold (Supplementary File 1). Previously reported mutations in the capsid proteins of icosahedral viruses such as the P22 bacteriophage have only resulted in the formation of smaller icosahedra (petite capsids), tubular filaments (polyheads) or aberrant polymers.^(15)^ There are also relatively few examples of non-viral protein cages with tetrahedral symmetry. Examples of wild-type tetrahedral protein cages include the archaeal ferritin from *Archeoglobus fulgidus* and the bacterial Dps mini-ferritins, which adopt a tetrahedral rather than the typical octahedral symmetry.^(33, 34)^ Engineered versions of the icosahedral lumazine synthase compartment from *Aquifex aeolicus* are some of the only mutant cages that have been observed to form tetrahedral symmetry.^(35)^ Like the mutant tetrahedral encapsulins, these lumazine synthase cages also have increased porosity due to sub-optimal packing of protomers but have a largely spherical morphology, lacking the distinct tetrahedral shape and flat triangular faces present in the mutant encapsulin 3×His-MxEnc. In these morphological terms, the mutant encapsulin is somewhat more reminiscent of *de novo* protein cages that are designed to assemble with tetrahedral symmetry.^(36–41)^

Another unusual property of the mutant encapsulins is their dimorphism, simultaneously forming stable icosahedral and tetrahedral assemblies from only one protomer sequence. There are limited prior reports of well-defined capsid polymorphs that co-exist under the same experimental conditions.^(13)^ The wild-type capsid protein of the beak and feather disease virus has been shown to form a D_5_-symmetric dimer of pentamers alongside the more common *T*=1 icosahedral virus-like particles.^(42)^ Meanwhile, a loop insertion mutant of the icosahedral MS2 bacteriophage capsid has been reported to generate extended nanotubes^(43)^ and a collection of low abundance *T*=4 icosahedral and symmetry-broken spherical polymorphs (D_5_, D_3-A_, D_3-B_ symmetry)^(44)^ alongside the predominant *T*=3 icosahedral form. These uncommon examples of polymorphism are likely to represent kinetic trapping of misassembled states along the pathways towards native capsid assembly.

The existence of tetrahedral encapsulin cages deepens our current understanding of capsid evolution and the potential differences between viral capsids and non-viral protein cages. The HK97 bacteriophage capsid and related capsids with the same protomeric fold have now been studied for well over 30 years, encompassing a viral family which includes some of the earliest studied viruses such as lambda phage.^(45)^ As icosahedral symmetry is well established as a ubiquitous requirement across this family of viruses,^(12, 46)^ the formation of tetrahedra represents an unexpected discovery. It may be that non-viral encapsulins are subject to fewer geometric constraints due to diminished evolutionary pressure for maintaining maximal volume and stability, as the contents of the cage are small enzymes rather than large nucleic acids. It remains to be seen whether similar mutations in other capsid proteins with the HK97 fold can result in tetrahedral capsids, or if indeed this polymorphism is unique to encapsulins and their cargo packaging mechanism.

The ability for a single point mutation to induce large-scale reorganisation of protomers has potential broader implications for protein cage design and the underlying nature of protein oligomerisation. The serendipitous formation of a novel protein-protein interaction in 3×His-MxEnc that drives cage assembly implies that the creation of novel cages with unique morphologies may occur more readily than expected. It is possible that a small change can propagate through an entire homomeric structure with only a small energetic driving force that is multiplied throughout the assembly to achieve a stable structure. This effect may account for the many early successes of designing cage-like structures,^(47)^ and the rapidly increasing success rates of *de novo* design with modern techniques such as machine learning.^(48)^ More generally, protein oligomerisation may occur more easily than intuitively expected, with small evolutionary changes sufficient to achieve the diversity of multimeric structures that exist across the set of all known proteins.^(49, 50)^

Finally, access to novel capsid symmetries opens new opportunities for technologies that depend on capsid morphology. One such example is the use of engineered capsids as scaffolds for vaccine development,^(10, 11)^ where the overall morphology can affect the number and positioning of antigens on the exterior surface, thus impacting efficacy. Different capsid morphologies and their resulting changes in porosity may also affect the efficiency of capsids that are engineered to house enzymes for biomanufacturing applications.^(26, 27)^ Thus, further study of how capsid geometry can be controlled is expected to complement *de novo* design efforts in generating a new suite of capsid-based tools with diverse properties for biotechnological applications.

## Materials and methods

### Molecular cloning

Plasmid constructs were cloned by Gibson assembly^(51)^ using NEBuilder HiFi DNA Assembly Master Mix (New England Biolabs). Codon-optimised gene fragments for wild-type MxEnc and the triple mutants 3×His-MxEnc, 3×Ala-MxEnc, 3×Asp-MxEnc, and 3×Lys-MxEnc were purchased as gBlocks (Integrated DNA Technologies) and cloned into a pETDuet-1 vector that was linearised with NdeI and AvrII restriction enzymes. Single mutants were cloned by Gibson assembly with the same linearised vector and two PCR products, covering the 5’ and 3’ segments of the gene with an overlapping region containing the desired mutation. Following Gibson assembly, plasmids were transformed into chemically competent DH5α *E. coli* cells. Sequences of all plasmids were confirmed by Sanger sequencing at the Australian Genome Research Facility. Gene fragment and primer sequences are provided in SI Section 1.

### Recombinant protein production

Plasmids were transformed and expressed in chemically competent BL21(DE3) *E. coli* cells. Overnight 10 mL cultures were diluted into 400 mL to obtain an OD_600_ of 0.05, then grown at 37 °C until an OD_600_ of 0.4-0.6 was reached. Protein expression was induced with 0.1 mM isopropyl β-D-1-thiogalactopyranoside (IPTG) and the culture was shaken at 18 °C overnight. Cells were then pelleted by centrifugation at 3900 × *g* for 20 min (5810R centrifuge with S-4-104 rotor, Eppendorf) and stored at −20 °C.

Frozen pellets were resuspended in 25 mL of lysis buffer (50 mM Tris pH 8, 200 mM NaCl, DNase I (10 μg/mL), lysozyme (100 μg/mL), 1× protease inhibitor (cOmplete Protease Inhibitor Cocktail, Roche). After keeping the suspension on ice for 30 min, cells were lysed using a Sonopuls HD 4050 probe sonicator with a TS-106 probe (Bandelin) at an amplitude setting of 55% and a pulse time of 8 s on and 10 s off for a total of 11-17 min. The lysate was clarified by centrifugation at 12,000 × *g* for 40 min. The supernatant was isolated and solid ammonium sulfate was added to a concentration of 20% (w/v). The sample was mixed on a rocker at 4 °C for 15 min before centrifugation at 10,000 × *g* for 15 min. The supernatant was isolated and a second portion of solid ammonium sulfate was added to achieve a final concentration of 40% (w/v). After rocking at 4 °C for another 15 min and centrifugation at 10,000 × *g* for 15 min, the supernatant was discarded and the pellet was resuspended in 5 mL of purification buffer (50 mM Tris pH 8, 200 mM NaCl). The sample was dialysed in 1 L of purification buffer at 4°C using Snakeskin Dialysis Tubing (3.5 k MWCO, Thermo Fisher), followed by an overnight dialysis under the same conditions.

The dialysed sample was filtered and purified over three chromatographic steps on an AKTA Pure or AKTA Start (Cytiva). Nucleic acids were removed by anion-exchange chromatography on a HiPrep Q XL 16/10 column at a flow rate of 5 mL/min. The encapsulin that eluted in the unretained peak was then purified by size-exclusion chromatography on a HiPrep 16/60 Sephacryl S-500 HR column at a flow rate of 1 mL/min. Fractions containing encapsulin were concentrated using Amicon Ultra-15 centrifugal filters (100 kDa MWCO, Merck), followed by a second round of size-exclusion chromatography on a Superose 6 Increase 10/300 GL column at a flow rate of 0.5 mL/min. The ‘early’ and ‘late’ fractions refer to elution at 11.5-12 mL and 13.2-13.7 mL, respectively. For population analysis, all fractions containing encapsulin were pooled for data collection. Purified encapsulins were quantified *via* their absorbance at 280 nm and stored at 4 °C until further use. Cargo loading of purified encapsulins filled with mNeonTP was estimated by measuring the absorbance at both 280 nm and 506 nm, using an extinction coefficient of 116000 M^-1^ cm^-1^.

### Mass photometry

Mass photometry was carried out on a TwoMP using Acquire MP and Discover MP software (Refeyn), using bovine serum albumin (66 kDa), apoferritin (480 kDa), and bovine thyroglobulin (670 kDa) as mass calibrants. Encapsulin samples were prepared at a concentration of 1 μM in 50 mM Tris pH 8 with 200 mM NaCl. One drop of buffer (∼10-18 μL) was applied to the gasket and focus was obtained using the Droplet-Dilution function. Subsequently, 2-10 μL of encapsulin solution was mixed with the buffer droplet and a 1 min recording was obtained. The final total sample volume for each recording was 20 μL, with a final encapsulin concentration in the 50-250 nM range.

### Blue native polyacrylamide gel electrophoresis

Blue native PAGE analysis was carried out using NativePAGE 3-12% Bis-Tris 1.0 mm Mini Protein Gels (Thermo Fisher). Samples contained 10 μg of protein in NativePAGE Sample Buffer. Anode buffer, dark blue cathode buffer and light blue cathode buffer were all prepared according to manufacturer protocols. Gel electrophoresis began with dark blue cathode buffer at 150 V for 5 min, followed by light blue cathode buffer for a further 75 min. Gels were then stained with Coomassie Brilliant Blue G (80 mg dissolved in 1 L water with 30 mM HCl) and destained with water before visualising using a ChemiDoc MP Imaging System (Bio-Rad). For mNeonTP-loaded encapsulins, gels were imaged using the epi-blue illumination source (460-490 nm) prior to staining.

### sNative charge detection mass spectrometry

CD-MS measurements were performed on homebuilt instruments as previously described.^(52–55)^ Briefly, samples were transferred into 200 mM ammonium acetate, a compatible MS salt, using Micro Bio-Spin P-6 gel columns before ion generation *via* a 200 nm diameter pulled quartz emitter (Sutter p2000) for nanoelectrospray ionisation. Ions entered the instrument through a metal capillary where they were thermalised and desolvated through a FUNPET interface. Upon exiting a series of ion optics, ions were focused into a dual hemispherical deflection energy analyser (HDA) for energy filtration. Ions with the set energy (130 eV/z) enter a detection cylinder housed within an electrostatic linear ion trap. Trapping times were either 400 ms with a charge uncertainty of ∼0.5 e or 1500 ms for quantised charge and an expected mass resolution of ∼330 Da. Each dataset contained a minimum of 8000 ions. Masses and standard deviations are reported based on Gaussian fitting of the data.

### Negative stain transmission electron microscopy

Gold grids (200-mesh coated with Formvar-carbon film, EMS) were made hydrophilic by glow discharge at 50 mA for 30 s or 5 mA for 60 s. The grid was floated on 20 µL of sample (100 µg/mL) for 1 min, blotted with filter paper, and washed once with 20 µL distilled water before staining with 20 µL of 2% (w/v) uranyl acetate solution or 0.75% (w/v) uranyl formate solution for 1 min. TEM images were captured at 120 keV using a JEM-1400 (JEOL) electron microscope at Sydney Microscopy and Microanalysis or at 100 keV using a Morgagni transmission electron microscope at the University of Michigan Life Sciences Institute.

### Sample preparation and Cryo-EM data collection

Grids of 3×His-MxEnc pore mutants were prepared by applying 3.5 µL of freshly prepared protein at 3.0 mg/mL to glow-discharged Quantifoil R2/1, 200 mesh copper holey carbon grids (EMS, Q225CR1). The grids were then frozen by plunging into liquid ethane using an FEI Vitrobot Mark IV with the following parameters: temperature 22 °C, humidity 100%, blot force 5, blot time 2 s. The grids were clipped and stored in liquid nitrogen until data collection.

Data was collected using an FEI Titan Krios G3 cryo-transmission electron microscope operating at 300 kV and equipped with a Gatan K3 Direct Detector with Bioquantum Imaging Filter. SerialEM^(56)^ was used to select targets and acquire movies with the following settings: defocus range -1 to -2.5 µm, dose of 50 e^-^/Å^2^, magnification 105,000×, exposure time 2.0972 s with 41.94 ms per frame. 1,475 movies were collected for the T1 dataset, and 3,765 movies were collected for the tetrahedral dataset, from individual grids. Further data collection information can be found in Table S5.1.

### Cryo-EM data processing and model building

All data processing was carried out using CryoSPARC v4.2.1.^(57)^ For the T1 dataset, 1,475 movies were motion corrected using patch motion correction, followed by patch CTF estimation. Movies with CTF fits worse than 5 Å were removed, resulting in 1,372 remaining movies. 509 particles were picked manually and used to generate initial templates by 2D classification. Template picker was then used to pick particles from all movies using a particle size of 250 Å, resulting in 140,872 extracted particles with a box size of 384 pixels. The particles were then sorted by two rounds of 2D classification resulting in 85,391 particles of T1 shells. The selected particles were then used to generate two ab-initio volumes with I symmetry imposed, with 85,150 particles contained in the main volume. Homogeneous refinement of the particles against the major ab-initio class with I symmetry, per-particle defocus refinement optimization, and per-group CTF parameter optimization, and Ewald sphere correction resulted in a 2.48 Å reconstruction. Heterogeneous refinement with three classes and I symmetry imposed was performed against the volume from the homogeneous refinement, resulting in a major class of particles containing 77,769 particles. These particles were then used to perform a homogeneous refinement with I symmetry imposed, per-particle defocus optimization, per-group CTF parameter optimization, and Ewald sphere correction applied, resulting in a final map with 2.43 Å resolution.

For the tetrahedral dataset, 3,765 movies were motion corrected using patch motion correction, followed by patch CTF estimation. Movies with CTF fits worse than 5 Å were removed, resulting in 3,691 movies. 521 particles were picked manually and used to generate initial templates by 2D classification. Template picker was then used to pick particles from all movies using a particle size of 200 Å, resulting in 495,482 extracted particles with a box size of 384 pixels. The particles were then sorted by two rounds of 2D classification resulting in 261,001 particles containing tetrahedral shells. The selected particles were then used to generate three ab-initio volumes with T symmetry imposed, with 260,795 particles contained in the main volume. These particles were sorted by heterogeneous refinement with T symmetry applied, resulting in a majority class containing 170,719 particles. Homogeneous refinement of these particles against the major heterogeneous refinement class volume with T symmetry, per-particle defocus refinement optimization, and per-group CTF parameter optimization, resulted in a 2.85 Å resolution reconstruction. To further improve the quality of the map, particle sets from the heterogeneous refinement containing 170,719 and 83,949 particles were then sorted by an additional round of heterogeneous refinement for each of these particle sets. 233,299 particles from the second round of heterogeneous refinement were symmetry expanded using T symmetry resulting in 2,799,588 particles. The symmetry-expanded particles were used as an input for a local refinement against the 2.85 Å resolution map with a single asymmetric unit (ASU) contained within the custom refinement mask. The following parameters were applied for the local refinement: use pose/shift gaussian prior during alignment, 7° standard deviation of prior over rotation, 4 Å standard deviation of prior over shifts, 0.05° maximum alignment resolution, re-center rotations each iteration, and re-center shifts each iteration. This local refinement resulted in an improved final map of the asymmetric unit with a resolution of 2.71 Å that was then used for model building.

To build the 3×His-MxEnc models, ESMfold^(58)^ was used to provide initial structures, which were then docked into the cryo-EM maps using ChimeraX v1.25^(59)^ by using the fit-to-volume command. The placed coordinates were then manually refined against the map using Coot v0.9.8.1^(60)^ until the fit was satisfactory. Phenix v1.20.1-4487^(61)^ was used to find the symmetry operators from the map of the T=1 shell using the symmetry-from-map function. The symmetry operators were then applied to the docked promoter model to generate a T=1 icosahedral shell containing 60 copies of the 3×His-MxEnc protomer. The model was refined using real-space refinement with NCS restraints, three macro cycles, rotamers fit outliers with target fix_outliers and tuneup outliers, and all other settings left to default. The model was further improved by iterative manual refinements using Coot. The final model was validated against the map using the comprehensive validation tool in Phenix to ensure that the model’s geometry and map fit were satisfactory.

The same model building and validation strategy was used for the tetrahedral model, except three copies of 3×His-MxEnc were placed in the ASU and real-space refinement was performed against the locally refined map without NCS restraints. The NCS matrices for the tetrahedral shell were obtained using the symmetry from map function in Phenix using the initial 2.85 Å resolution map with T symmetry imposed as an input.

The models and cryo-EM densities for the T1 and tetrahedral 3×His-MxEnc assemblies were deposited in the Protein Data Bank (PDB) and the Electron Microscopy Data Bank (EMDB) under the PDB IDs 8V4N, and 8V4Q, and EMDB IDs EMD-42974, and EMD-42975, respectively.

## Supporting information

PDB Tetrahedral Cages List

## Acknowledgements and funding sources

YHL acknowledges funding from the Australian Research Council (DE19010062, DP230101045) and Westpac Scholars Trust (WRF2020). We acknowledge the core facilities at Sydney Analytical and Sydney Microscopy and Microanalysis for providing infrastructure support. TNS, FL, LSRA, and YHL acknowledge support from the ARC Centre of Excellence in Synthetic Biology, while YHL also acknowledges support from the ARC Centre of Excellence for Innovations in Peptide and Protein Science. TWG acknowledges funding from the NIH (R35GM133325). Research reported in this publication was supported by the University of Michigan Cryo-EM Facility (U-M Cryo-EM). U-M Cryo-EM is grateful for support from the U-M Life Sciences Institute and the U-M Biosciences Initiative. Molecular graphics and analyses performed with UCSF ChimeraX, developed by the Resource for Biocomputing, Visualization, and Informatics at the University of California, San Francisco, with support from the National Institutes of Health R01GM129325 and the Office of Cyber Infrastructure and Computational Biology, National Institute of Allergy and Infectious Diseases. BED and MFJ are shareholders in Megadalton Solutions, a company that is engaged in commercializing CD-MS. BED is an employee of Megadalton Solutions and MFJ is a consultant for Waters. RT acknowledges the Wellcome Trust for financial support through the Joint Investigator Award (110145 & 110146), the EPSRC for an Established Career Fellowship (EP/R023204/1) which also provided funding for FF, and the Royal Society for a Royal Society Wolfson Fellowship (RSWF/R1/180009).

