## Supporting Information for "Point mutation in a virus-like capsid drives symmetry reduction to form tetrahedral cages"

**Contents**

1. Gene fragment and primer sequences
2. SDS-PAGE gels during encapsulin purification
3. Supplementary native PAGE gels
4. Supplementary mass photometry data
5. Supplementary cryo-electron microscopy data
6. Supplementary charge detection mass spectrometry data
7. ConSurf analysis of amino acid conservation
8. Planar embedding of surfaces into a gyrated lattice

### 1. Gene fragment and primer sequences

#### 1.1 Codon-optimised sequence of wild-type MxEnc gene

ATGCCGGATTTCTTGGGACATGCTGAGAACCCATTGCGCGAAGAAGAGTGGGCCCGTTTAAACGAGAC
TGTAATCCAGGTTGCCCGCCGCTCTTTGGTTGGGCGTCGTATCTTAGACATTTATGGCCCTTTGGGAG
CGGGTGTGCAAACGGTTCCATATGATGAGTTTCAGGGCGTAAGTCCGGGGGCCGTTGATATCGTGGGT
GAGCAAGAAACCGCAATGGTGTTTACCGATGCCCGCAAGTTTAAGACCATCCCGATCATTTATAAAGA
TTTCTTGCTGCATTGGCGTGATATTGAGGCGGCGCGTACCCATAACATGCCCTTAGATGTTTCGGCAG
CCGCTGGGGCGGCGGCCTTATGCGCTCAGCAGGAAGACGAGTTAATCTTCTACGGCGACGCTCGCTTG
GGATACGAAGGGTTAATGACAGCTAATGGCCGTCTGACGGTTCCTTTAGGGGACTGGACCTCTCCGGG
GGGTGGTTTTCAGGCAATCGTGGAAGCTACCCGCAAACTGAACGAGCAGGGACATTTTGGTCCCTACG
CCGTGGTTTTATCGCCTCGCTTGTATTCCCAACTTCACCGCATCTACGAAAAAACAGGGGTGCTGGAA
ATTGAAACGATTCGTCAACTTGCCTCTGATGGTGTGTATCAATCAAACCGTCTGCGCGGGGAATCTGG
TGTTGTGGTGTCCACAGGCCGTGAAAATATGGACTTAGCGGTAAGCATGGACATGGTCGCTGCTTATT
TAGGAGCTTCTCGTATGAATCACCCTTTTCGCGTACTGGAAGCATTGCTTCTGCGCATTAAGCACCCA
GATGCGATTTGTACATTAGAAGGAGCTGGAGCTACAGAGCGCCGCTAA

#### 1.2 Codon-optimised sequence of mNeonGreen with targeting peptide for encapsulation

The mNeonGreen expression construct was designed with an *N*-terminal hexahistidine tag and a *C*-terminal targeting peptide sequence from *M. xanthus*, using the second cassette of pCDFDuet-1 plasmid (NdeI and PacI restriction sites).

ATGCACCACCATCATCATCACGGAGGTTCAGTCTCCAAGGGAGAGGAGGACAATATGGCTAGTCTTCC
GGCCACTCATGAGTTACATATCTTCGGATCCATAAACGGCGTTGATTTCGATATGGTGGGTCAAGGCA
CTGGTAACCCCAATGATGGCTACGAAGAACTTAACTTAAAATCTACTAAAGGCGATCTTCAGTTTTCC
CCATGGATACTTGTGCCTCATATTGGCTACGGGTTTCACCAATATCTGCCTTATCCGGATGGAATGTC
CCCCTTCCAAGCCGCAATGGTAGACGGCAGTGGCTATCAAGTCCACCGTACCATGCAGTTTGAAGATG
GCGCATCCCTGACAGTTAATTATCGGTATACATACGAGGGGTCGCATATTAAAGGAGAGGCGCAAGTC
AAGGGGACAGGGTTTCCCGCCGATGGGCCAGTCATGACAAACTCGTTAACTGCCGCCGACTGGTGCAG
ATCGAAGAAAACCTACCCAAACGATAAGACGATCATATCTACCTTTAAATGGTCTTACACTACGGGTA
ACGGAAAACGCTACAGATCAACCGCGCGGACAACGTACACCTTTGCTAAGCCCATGGCAGCGAACTAC
TTGAAGAATCAGCCGATGTACGTGTTTAGAAAGACCGAGCTTAAACACTCGAAAACTGAATTGAATTT
TAAAGAATGGCAGAAAGCTTTTACGGACGTAATGGGCATGGACGAACTGTATAAGTCAGGCGGGCCAT
TAACGGTTGGATCGCTGCGTCGTGGAGGCTAA

#### 1.3 Gene fragments for Gibson assembly of triple mutants

Coding regions of gene fragments are shown in capital letters, while homology for Gibson assembly is shown in lower case letters. Mutant region is shown in bold letters.

##### 3×His-MxEnc

gttaagtataagaaggagatatacatATGCCGGATTTCTTGGGACATGCTGAGAACCCATTGCGCGAA
GAAGAGTGGGCCCGTTTAAACGAGACTGTAATCCAGGTTGCCCGCCGCTCTTTGGTTGGGCGTCGTAT
CTTAGACATTTATGGCCCTTTGGGAGCGGGTGTGCAAACGGTTCCATATGATGAGTTTCAGGGCGTAA
GTCCGGGGGCCGTTGATATCGTGGGTGAGCAAGAAACCGCAATGGTGTTTACCGATGCCCGCAAGTTT
AAGACCATCCCGATCATTTATAAAGATTTCTTGCTGCATTGGCGTGATATTGAGGCGGCGCGTACCCA
TAACATGCCCTTAGATGTTTCGGCAGCCGCTGGGGCGGCGGCCTTATGCGCTCAGCAGGAAGACGAGT
TAATCTTCTACGGCGACGCTCGCTTGGGATACGAAGGGTTAATGACAGCTAATGGCCGTCTGACGGTT
CCTTTAGGGGACTGGACCTCTCCGGGTGGTGGTTTTCAGGCAATCGTGGAAGCTACCCGCAAACTGAA
CGAGCAGGGACATTTTGGTCCCTACGCCGTGGTTTTATCGCCTCGCTTGTATTCCCAACTTCACCGCA
TCTACGAA**CATCATCAT**GTGCTGGAAATTGAAACGATTCGTCAACTTGCCTCTGATGGTGTGTATCAA
TCAAACCGTCTGCGCGGGGAATCTGGTGTTGTGGTGTCCACAGGCCGTGAAAATATGGACTTAGCGGT
AAGCATGGACATGGTCGCTGCTTATTTAGGAGCTTCTCGTATGAATCACCCTTTTCGCGTACTGGAAG
CATTGCTTCTGCGCATTAAGCACCCAGATGCGATTTGTACATTAGAAGGAGCTGGAGCTACAGAGCGC
CGCTAAcctaggctgctgccaccgctg

##### 3×Ala-MxEnc

gttaagtataagaaggagatatacatATGCCGGATTTCTTGGGACATGCTGAGAACCCATTGCGCGAA
GAAGAGTGGGCCCGTTTAAACGAGACTGTAATCCAGGTTGCCCGCCGCTCTTTGGTTGGGCGTCGTAT
CTTAGACATTTATGGCCCTTTGGGAGCGGGTGTGCAAACGGTTCCATATGATGAGTTTCAGGGCGTAA
GTCCGGGGGCCGTTGATATCGTGGGTGAGCAAGAAACCGCAATGGTGTTTACCGATGCCCGCAAGTTT
AAGACCATCCCGATCATTTATAAAGATTTCTTGCTGCATTGGCGTGATATTGAGGCGGCGCGTACCCA
TAACATGCCCTTAGATGTTTCGGCAGCCGCTGGGGCGGCGGCCTTATGCGCTCAGCAGGAAGACGAGT
TAATCTTCTACGGCGACGCTCGCTTGGGATACGAAGGGTTAATGACAGCTAATGGCCGTCTGACGGTT
CCTTTAGGGGACTGGACCTCTCCGGGTGGTGGTTTTCAGGCAATCGTGGAAGCTACCCGCAAACTGAA
CGAGCAGGGACATTTTGGTCCCTACGCCGTGGTTTTATCGCCTCGCTTGTATTCCCAACTTCACCGCA
TCTACGAA**GCCGCAGCG**GTGCTGGAAATTGAAACGATTCGTCAACTTGCCTCTGATGGTGTGTATCAA
TCAAACCGTCTGCGCGGGGAATCTGGTGTTGTGGTGTCCACAGGCCGTGAAAATATGGACTTAGCGGT
AAGCATGGACATGGTCGCTGCTTATTTAGGAGCTTCTCGTATGAATCACCCTTTTCGCGTACTGGAAG
CATTGCTTCTGCGCATTAAGCACCCAGATGCGATTTGTACATTAGAAGGAGCTGGAGCTACAGAGCGC
CGCTAAcctaggctgctgccaccgctg

**3×Asp-MxEnc**

gttaagtataagaaggagatatacatATGCCGGATTTCTTGGGACATGCTGAGAACCCATTGCGCGAA
GAAGAGTGGGCCCGTTTAAACGAGACTGTAATCCAGGTTGCCCGCCGCTCTTTGGTTGGGCGTCGTAT
CTTAGACATTTATGGCCCTTTGGGAGCGGGTGTGCAAACGGTTCCATATGATGAGTTTCAGGGCGTAA
GTCCGGGGGCCGTTGATATCGTGGGTGAGCAAGAAACCGCAATGGTGTTTACCGATGCCCGCAAGTTT
AAGACCATCCCGATCATTTATAAAGATTTCTTGCTGCATTGGCGTGATATTGAGGCGGCGCGTACCCA
TAACATGCCCTTAGATGTTTCGGCAGCCGCTGGGGCGGCGGCCTTATGCGCTCAGCAGGAAGACGAGT
TAATCTTCTACGGCGACGCTCGCTTGGGATACGAAGGGTTAATGACAGCTAATGGCCGTCTGACGGTT
CCTTTAGGGGACTGGACCTCTCCGGGTGGTGGTTTTCAGGCAATCGTGGAAGCTACCCGCAAACTGAA
CGAGCAGGGACATTTTGGTCCCTACGCCGTGGTTTTATCGCCTCGCTTGTATTCCCAACTTCACCGCA
TCTACGAA**GATGACGAT**GTGCTGGAAATTGAAACGATTCGTCAACTTGCCTCTGATGGTGTGTATCAA
TCAAACCGTCTGCGCGGGGAATCTGGTGTTGTGGTGTCCACAGGCCGTGAAAATATGGACTTAGCGGT
AAGCATGGACATGGTCGCTGCTTATTTAGGAGCTTCTCGTATGAATCACCCTTTTCGCGTACTGGAAG
CATTGCTTCTGCGCATTAAGCACCCAGATGCGATTTGTACATTAGAAGGAGCTGGAGCTACAGAGCGC
CGCTAAcctaggctgctgccaccgctg

**3×Lys-MxEnc**

gttaagtataagaaggagatatacatATGCCGGATTTCTTGGGACATGCTGAGAACCCATTGCGCGAA
GAAGAGTGGGCCCGTTTAAACGAGACTGTAATCCAGGTTGCCCGCCGCTCTTTGGTTGGGCGTCGTAT
CTTAGACATTTATGGCCCTTTGGGAGCGGGTGTGCAAACGGTTCCATATGATGAGTTTCAGGGCGTAA
GTCCGGGGGCCGTTGATATCGTGGGTGAGCAAGAAACCGCAATGGTGTTTACCGATGCCCGCAAGTTT
AAGACCATCCCGATCATTTATAAAGATTTCTTGCTGCATTGGCGTGATATTGAGGCGGCGCGTACCCA
TAACATGCCCTTAGATGTTTCGGCAGCCGCTGGGGCGGCGGCCTTATGCGCTCAGCAGGAAGACGAGT
TAATCTTCTACGGCGACGCTCGCTTGGGATACGAAGGGTTAATGACAGCTAATGGCCGTCTGACGGTT
CCTTTAGGGGACTGGACCTCTCCGGGTGGTGGTTTTCAGGCAATCGTGGAAGCTACCCGCAAACTGAA
CGAGCAGGGACATTTTGGTCCCTACGCCGTGGTTTTATCGCCTCGCTTGTATTCCCAACTTCACCGCA
TCTACGAA**AAGAAGAAA**GTGCTGGAAATTGAAACGATTCGTCAACTTGCCTCTGATGGTGTGTATCAA
TCAAACCGTCTGCGCGGGGAATCTGGTGTTGTGGTGTCCACAGGCCGTGAAAATATGGACTTAGCGGT
AAGCATGGACATGGTCGCTGCTTATTTAGGAGCTTCTCGTATGAATCACCCTTTTCGCGTACTGGAAG
CATTGCTTCTGCGCATTAAGCACCCAGATGCGATTTGTACATTAGAAGGAGCTGGAGCTACAGAGCGC
CGCTAAcctaggctgctgccaccgctg

#### 1.4 Primers used for construction of single mutants

PCR was conducted with Q5 High-Fidelity 2× Master Mix (New England Biolabs) according to manufacturer protocols, using the wild-type MxEnc sequence from SI Section 1.1 as a template.

Homology regions between PCR fragments are underlined, with the codon change in bold. Lower case letter designate homology between PCR fragment and linearised pETDuet-1 vector.

**K199H_MxEnc**

PCR fragment 1:

MxEnc_F ggagatatacatATGCCGGATTTCTTGGGAC

K199H_R CAATTTC<u>CAGCACCCCTGT**ATG**TTCGTAGATGCG</u>

PCR fragment 2:

K199H_F: CAC<u>CGCATCTACGAA**CAT**ACAGGGGTGCTG</u>

MxEnc_R: cagcctaggTTAGCGGCGCTCTGTAG

**T200H_MxEnc**

PCR fragment 1:

MxEnc_F: ggagatatacatATGCCGGATTTCTTGGGAC

T200H_R: CAATTT<u>CCAGCACCCC**ATG**TTTTTCGTAGATGC</u>

PCR fragment 2:

T200H_F: CC<u>GCATCTACGAAAAA**CAT**GGGGTGCTGG</u>

MxEnc_R: cagcctaggTTAGCGGCGCTCTGTAG

**G201H:**

PCR fragment 1:

MxEnc_F: ggagatatacatATGCCGGATTTCTTGGGAC

G201H_R: <u>CAATTTCCAGCAC**ATG**TGTTTTTTCG</u>TAGATG

PCR fragment 2:

G201H_F: <u>CGAAAAAACA**CAT**GTGCTGGAAATTG</u>AAACGATTCG

MxEnc_R: cagcctaggTTAGCGGCGCTCTGTAG

### 2. SDS-PAGE gels during encapsulin purification

#### 2.1 Representative gels during the purification of 3×His-MxEnc

SDS-PAGE was conducted on Any kD Mini-Protean TGX Stain-Free protein gels (Bio-Rad) according to manufacturer protocols.

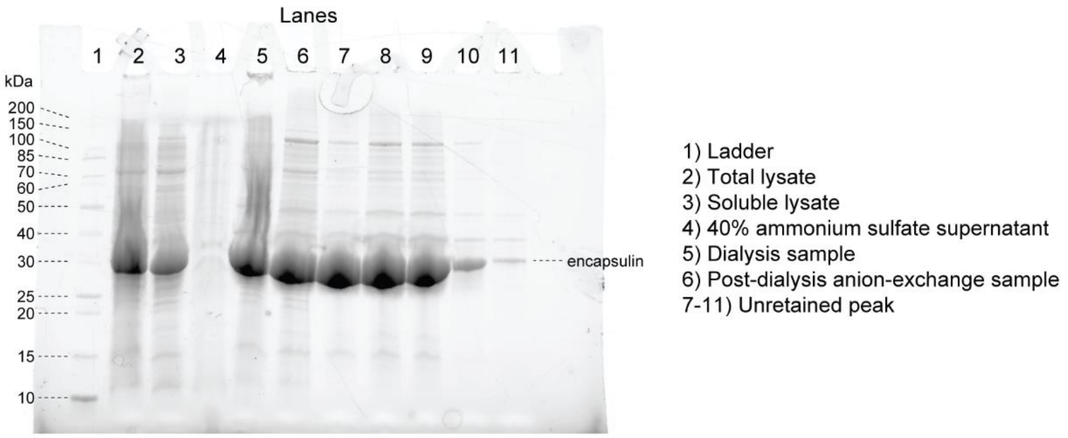

**Figure S2.1** SDS-PAGE gel of the early stages of purification for 3×His-MxEnc, including ammonium sulfate precipitation, dialysis, and anion-exchange chromatography on a HiPrep Q XL 16/60 column where the encapsulin elutes in the unretained peak (7.5-14 mL). The ladder is Unstained Protein Standard Broad Range (10-200 kDa, New England Biolabs).

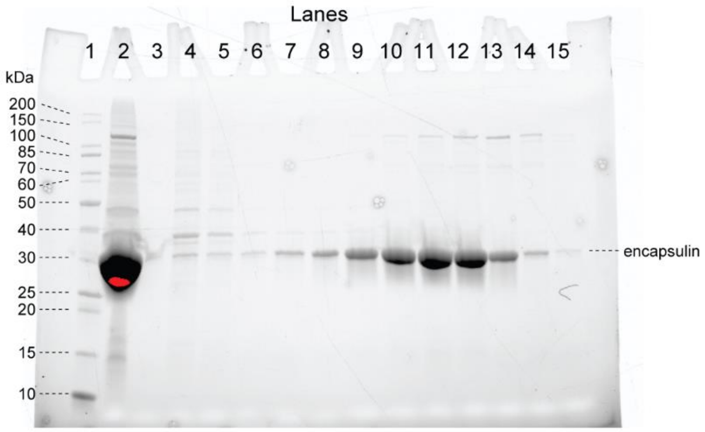

**Figure S2.2** SDS-PAGE gel of the low-resolution size-exclusion chromatography step for 3×His-MxEnc, conducted on a HiPrep 16/60 Sephacryl S-500 HR column. Lane 2 is the sample loaded for chromatography, while lanes 3-15 are the fractions (31.5-103 mL). The ladder in lane 1 is Unstained Protein Standard Broad Range (10-200 kDa, New England Biolabs). The size-exclusion chromatogram is shown in the manuscript in Figure 4a.

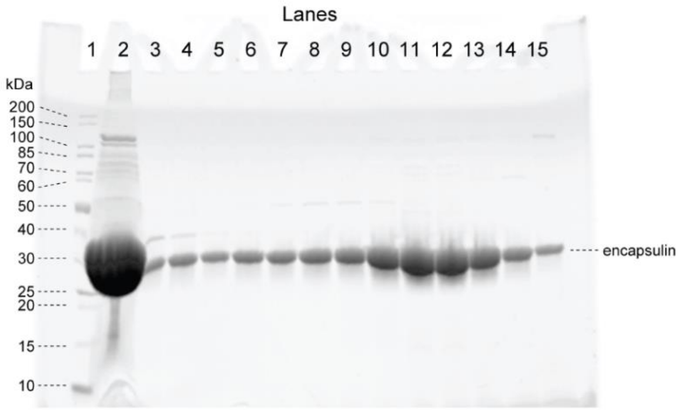

**Figure S2.3** SDS-PAGE gel of the high-resolution size-exclusion chromatography step for 3×His-MxEnc, conducted on a Superose 6 Increase 10/300 GL column. Lane 2 is the sample loaded for chromatography, while lanes 3-15 are the fractions (7.7-14.2 mL). The ladder in lane 1 is Unstained Protein Standard Broad Range (10-200 kDa, New England Biolabs). The size-exclusion chromatogram is shown in the manuscript in Figure 4b.

### 2.2 SDS-PAGE of 3×His-MxEnc co-expressed with mNeonTP

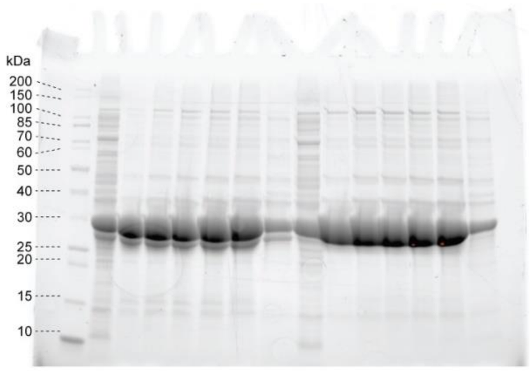

**Figure S2.4** SDS-PAGE gel of fractions from the anion-exchange chromatography step for 3×His-MxEnc + mNeonTP (left), in comparison to encapsulin expressed alone (right), conducted on a HiPrep Q XL 16/60 column. The lanes on the left of the gel show a clear additional band under the major encapsulin band, which is not present in the lanes on the right.

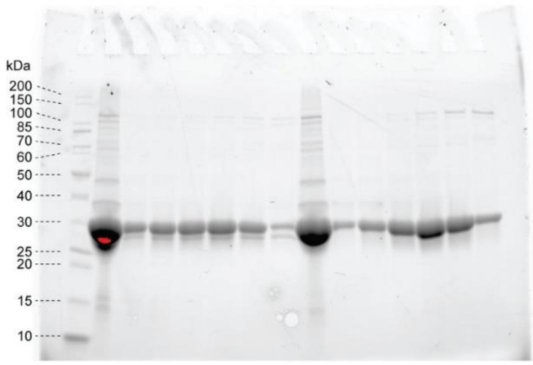

**Figure S2.5** SDS-PAGE gel of fractions from the low-resolution size-exclusion chromatography step for 3×His-MxEnc + mNeonTP (left), in comparison to encapsulin expressed alone (right), conducted on a HiPrep 16/60 Sephacryl S-500 HR column. The lanes on the left of the gel show a clear additional band under the major encapsulin band, which is not present in the lanes on the right.

### 3. Supplementary native PAGE gels

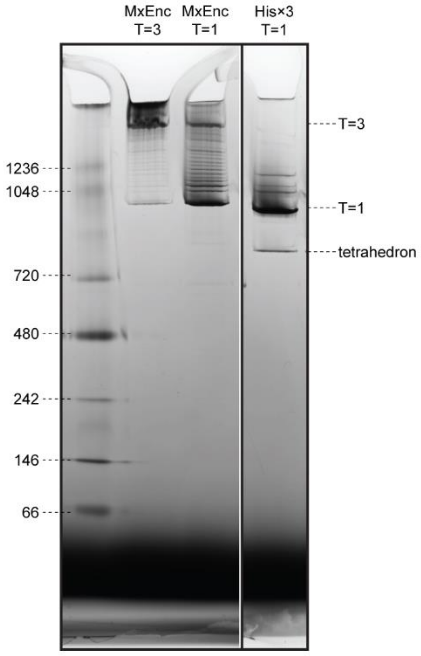

**Figure S3.1** Various samples from a blue native PAGE gel, showing the fractions from size-exclusion chromatography for wild-type MxEnc where the peaks for the T=3 and T=1 icosahedral assemblies were analysed. There is no evidence of a tetrahedral assembly in this sample, whereas the equivalent T=1 icosahedral peak for 3×His-MxEnc (run on a non-adjacent lane of the same gel) shows a clear lower band that corresponds to a tetrahedral assembly. The ladder used in the leftmost lane is the Native Mark Unstained Protein Standard (Thermo Fisher), which does not appear to run true to size relative to encapsulins under blue native conditions.

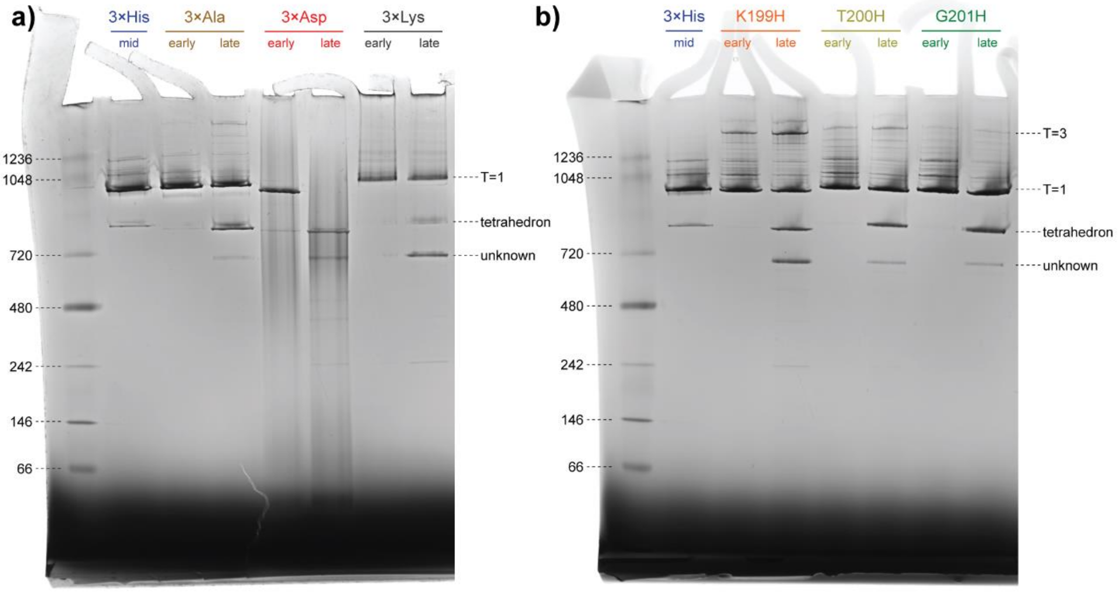

**Figure S3.2 a)** Blue native PAGE gel showing the full lanes for the data presented in the manuscript in Figure 4c. The streaking of 3×Asp-MxEnc is evident throughout the entire lane. The sample from the late fraction of 3×Lys-MxEnc shows an unidentified band of lower native mass, which may represent a smaller assembled state that we were unable to isolate for structural characterisation. **b)** Blue native PAGE gel showing the full lanes for the data presented in the manuscript in Figure 5b, where the potential smaller assembly is most prominent in the late fraction of the K199H-MxEnc sample (also see Figure S4.1).

### 4. Supplementary mass photometry data

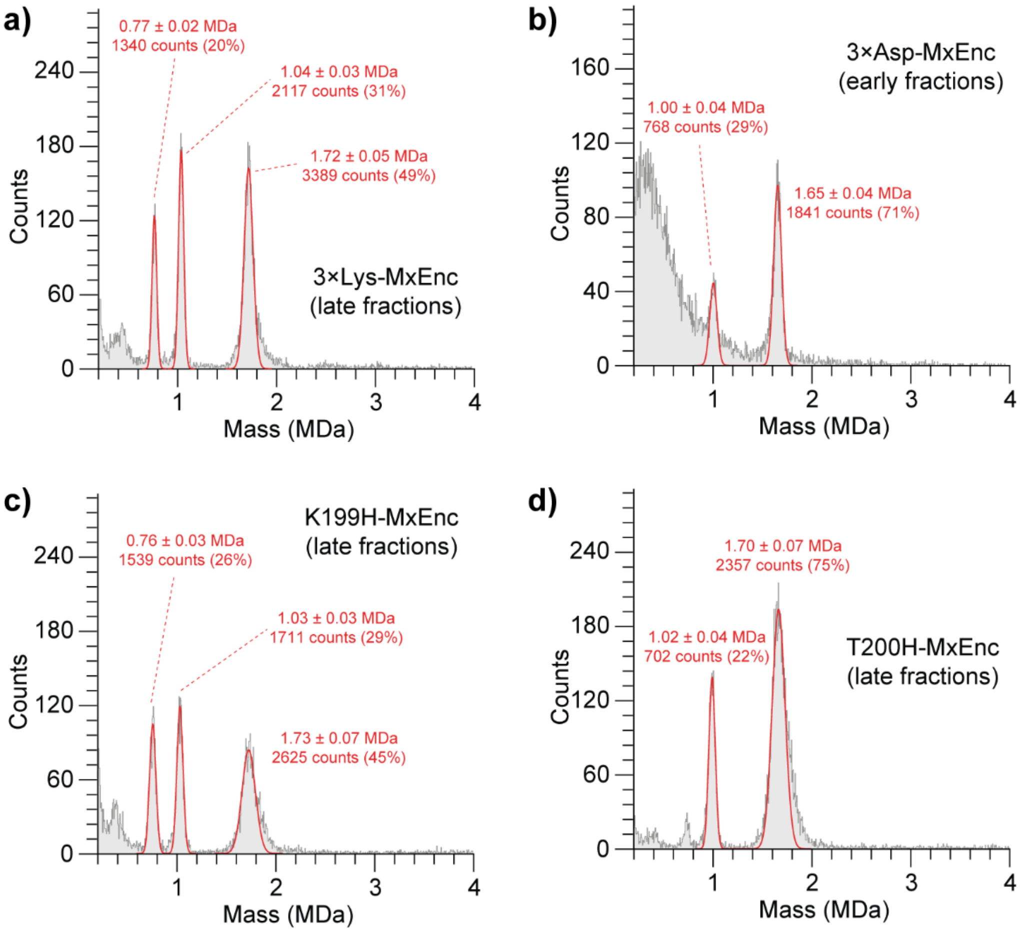

**Figure S4.1** Supplementary mass photometry data. **a)** 3×Lys-MxEnc shows evidence of tetrahedral and T=1 icosahedral forms, as well as an unidentified lower molecular weight species. **b)** 3×Asp-MxEnc shows evidence of tetrahedral and T=1 icosahedral forms. **c)** K199H-MxEnc shows evidence of tetrahedral and T=1 icosahedral forms, as well as an unidentified lower molecular weight species. **d)** T200H-MxEnc shows evidence of tetrahedral and T=1 icosahedral forms.

### 5. Supplementary cryo-electron microscopy data

#### 5.1 Cryo-EM data collection and model statistics

**Table S5.1** Data collection and model refinement statistics.

| | 3×His-MxEnc T1 (PDB ID: 8V4N, EMDB-42974) | 3×His-MxEnc Tetrahedron (PDB ID: 8V4Q, EMDB-42975) |
|---|---|---|
| **Data collection and processing** | | |
| Microscope | FEI Titan Krios G3 | FEI Titan Krios G3 |
| Detector | Gatan K3 | Gatan K3 |
| Magnification | 105,000x | 105,000x |
| Voltage (kV) | 300 | 300 |
| Electron exposure ($e^{-}$/Å$^2$) | 50 | 50 |
| Exposure time (s) | 2.097 | 2.097 |
| Frame time (ms) | 41.9 | 41.9 |
| Defocus range (μm) | -1.0 to -2.5 | -1.0 to -2.5 |
| Pixel size (Å) | 0.87 | 0.87 |
| Symmetry imposed | I | C1 (local refinement) |
| Final particle count (no.) | 77,769 | 2,799,588 |
| Map resolution (Å) | 2.43 | 2.71 |
| FSC threshold | 0.143 | 0.143 |
| **Refinement** | | |
| Initial model used | ESMfold | ESMfold |
| Model resolution (Å) | 2.7 | 2.8 |
| FSC threshold | 0.5 | 0.5 |
| Map sharpening *B* factor (Å$^2$) | -98.5 | -118.3 |
| Model composition | | |
| Non-hydrogen atoms | 2067 | 5993 |
| Protein residues | 263 | 761 |
| Chains | 1 | 3 |
| *B* factors (Å$^2$) | | |
| Protein | 47.46 | 38.93 |
| rms deviations | | |
| Bond lengths (Å) | 0.004 | 0.003 |
| Bond angles (°) | 0.940 | 0.525 |
| Validation | | |
| MolProbity score | 1.27 | 0.92 |
| Clashscore | 4.14 | 1.68 |
| Rotamer outliers (%) | 0.93 | 0.48 |
| Ramachandran plot | | |
| Favored (%) | 97.68 | 98.26 |
| Allowed (%) | 2.32 | 1.74 |
| Disallowed (%) | 0 | 0 |

### 5.2 Cryo-EM workflow figures

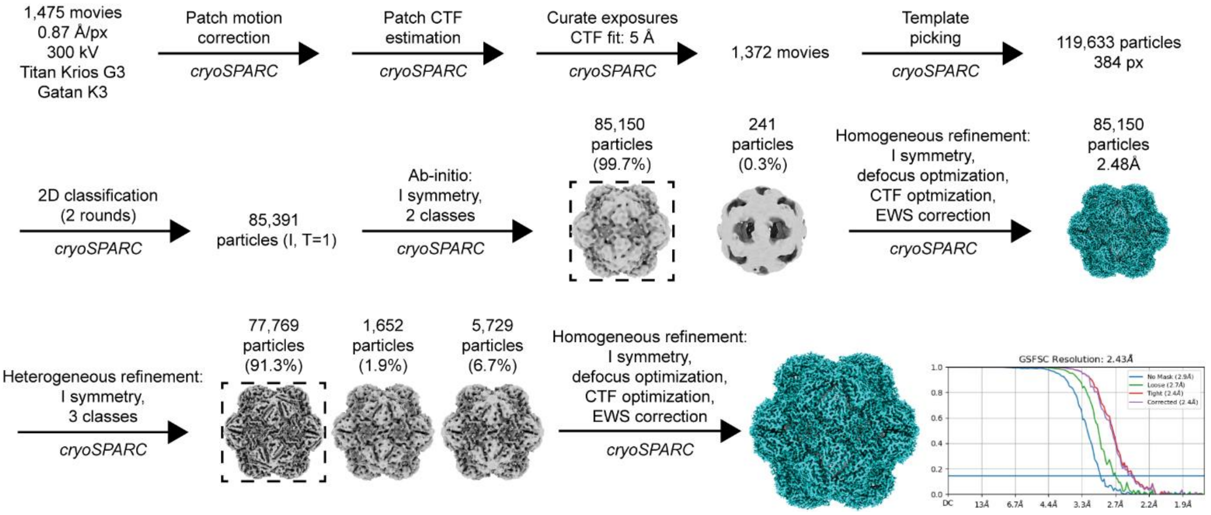

**Figure S5.1** Cryo-EM data processing workflow for the T=1 icosahedral assembly of 3×His-MxEnc.

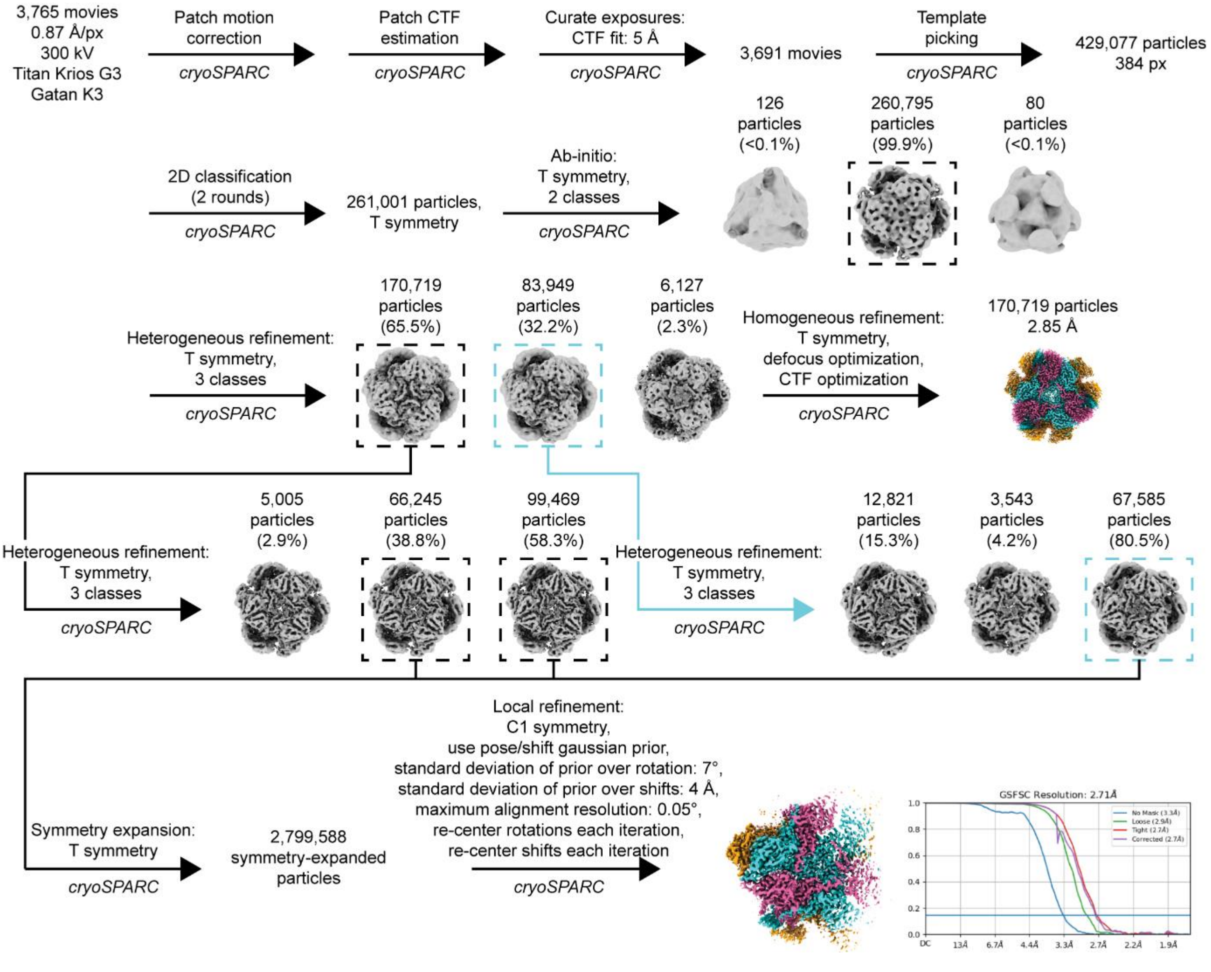

**Figure S5.2** Cryo-EM data processing workflow for the tetrahedral assembly of 3×His-MxEnc.

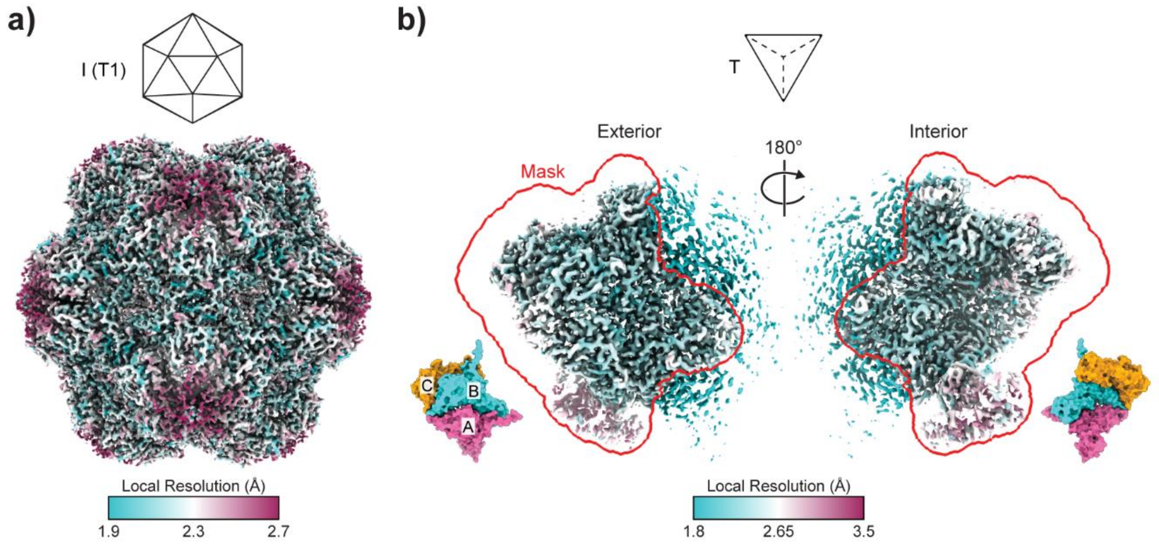

**Figure S5.3 Local resolution estimation of cryo-EM maps. a)** Estimation for the T=1 icosahedral assembly of 3×His-MxEnc based on an FSC threshold of 0.143. **b)** Estimation for the asymmetric unit (ASU) of the tetrahedral assembly of 3×His-MxEnc based on an FSC threshold of 0.143. The refinement mask used for masked local refinement is outlined in red. The orientation of the ASU map is indicated by an ASU model colored by unique ASU protomer (A: pink, B: cyan, C: gold).

### 6. Supplementary charge detection mass spectrometry data

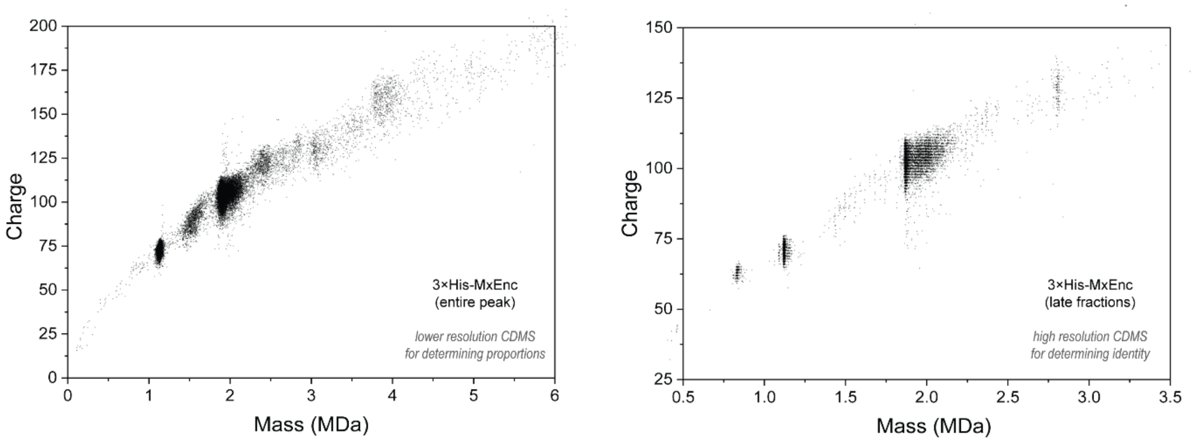

**Figure S6.1** Charge versus mass scatter plots that correspond to the native mass spectra shown in the manuscript in Figure 1d.

#### 7. ConSurf analysis of amino acid conservation

ConSurf analysis was conducted using the web server found at http://consurf.tau.ac.il, using PDB 7S20 Chain B as the input.

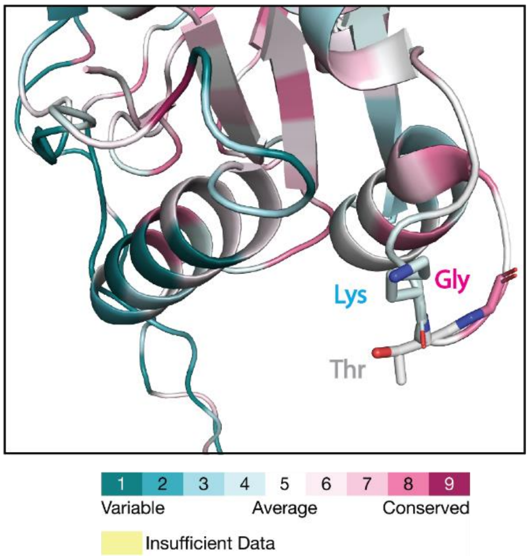

**Figure S7.1** ConSurf analysis of PDB 7S20 Chain B shows K199 is the least conserved, T200 is average, and G201 is the most conserved residue in the motif, suggesting that mutating G201 may potentially be detrimental to the formation of the native T=3 assembly.

### 8. Planar embedding of surfaces into a gyrated lattice

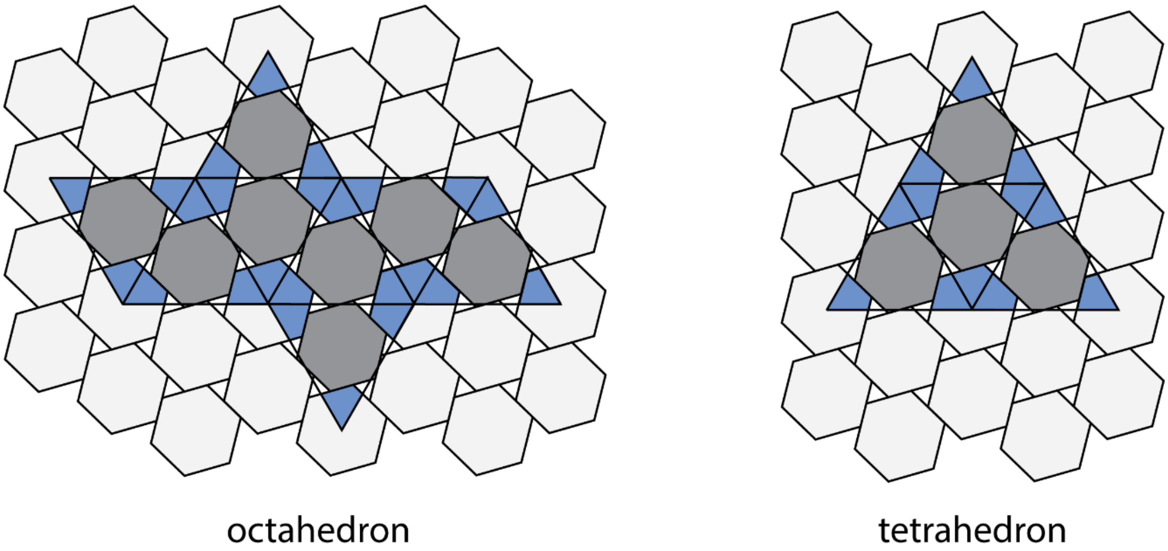

**Figure S8.1.** Planar embeddings of an octahedral and a tetrahedral surface into a gyrated hexagonal lattice. These geometric arrangements require the formation of encapsulin tetramers or trimers (blue) alongside the known hexamers (grey) in the final assembly. It is unlikely that a single type of protomer will be able to form these vastly different types of multimers.
