## Supplementary material for "Point mutation in a virus-like capsid drives symmetry reduction to form tetrahedral cages": PDB Tetrahedral Cages List

**SUPPLEMENTARY FILE 1**

**List of all PDB tetrahedral protein cages/containers/compartments** - no tetrahedral examples are viral capsids (or share the same fold as a viral capsid)

**PDB search date: 5th December 2023**

**PDB search terms:**
Full Text = "compartment"
Symmetry Type = "tetrahedral"

| Type (author annotated) | Identifier Entry ID | StructureData Structure Author | PubMed ID | DOI | Structure Title |
|---|---|---|---|---|---|
| Bacterial endoprotease | 3PV2 | Wrase, R., Scott, H., Hilgenfeld, R., Hansen, G. | 21670246 | 10.1073/pnas.1101084108 | Structure of Legionella fallonii DegQ (wt) |
| Bacterial endoprotease | 3PV5 | Wrase, R., Scott, H., Hilgenfeld, R., Hansen, G. | 21670246 | 10.1073/pnas.1101084108 | Structure of Legionella fallonii DegQ (N189G/P190G variant) |
| Bacterial endoprotease | 3PV3 | Wrase, R., Scott, H., Hilgenfeld, R., Hansen, G. | 21670246 | 10.1073/pnas.1101084108 | Structure of Legionella fallonii DegQ (S193A variant) |
| Animal ocular structural proteins | 2YGD | Braun, N., Zacharias, M., Peschek, J., Kastenmueller, A., Zou, J., Hanzlik, M., Haslbeck, M., Rappsilber, J., Buchner, J., Weinkauf, S. | 22143763 | 10.1073/pnas.1111014108 | Molecular architectures of the 24meric eye lens chaperone alphaB- crystallin elucidated by a triple hybrid approach |
| Bacterial dodecin (flavin storage) | 3J07 | Jehle, S., Vollmar, B., Bardiaux, B., Dove, K.K., Rajagopal, P., Gonen, T., Oschkinat, H., Klevit, R.E. | 21464278 | 10.1073/pnas.1014656108 | Model of a 24mer alphaB-crystallin multimer |
| Bacterial dodecin (flavin storage) | 2YJ0 | Vinzenz, X., Grosse, W., Linne, U., Meissner, B., Essen, L.-O. | 21897938 | 10.1039/c1cc12929e | X-ray structure of chemically engineered Mycobacterium tuberculosis Dodecin |
| Bacterial dodecin (flavin storage) | 2YIZ | Vinzenz, X., Grosse, W., Linne, U., Meissner, B., Essen, L.-O. | 21897938 | 10.1039/c1cc12929e | X-ray structure of Mycobacterium tuberculosis Dodecin |
| Amyloid | 5HOW | Kreutzer, A.G., Nowick, J.S., Spencer, R.K. | 26967810 | 10.1021/jacs.6b01332 | X-ray crystallographic structure of an Abeta 17-36 beta-hairpin. LV(PHI)FAEDCGSNKCAII(SAR)L(ORN)V |
| Amyloid | 5HOX | Kreutzer, A.G., Nowick, J.S., Spencer, R.K. | 26967810 | 10.1021/jacs.6b01332 | X-ray crystallographic structure of an A-beta 17_36 beta-hairpin. Synchrotron data set. (LVFFAEDCGSNKCAII(SAR)LMV). |
| Amyloid | 5HOY | Kreutzer, A.G., Spencer, R.K., Nowick, J.S. | 26967810 | 10.1021/jacs.6b01332 | X-ray crystallographic structure of an A-beta 17_36 beta-hairpin. X-ray diffractometer data set. (LVFFAEDCGSNKCAII(SAR)LMV). |
| Amyloid | 7JXO | Kreutzer, A.G., Haerianardakani, S., Nowick, J.S. | 33237748 | 10.1021/jacs.0c09281 | Triangular trimer of beta-hairpins derived from Abeta17-36 with an F20Cha mutation |
| Amyloid | 7U4P | Kreutzer, A.G., Haerianardakani, S., Nowick, J.S. | | | Covalently stabilized triangular trimer composed of Abeta17-36 beta-hairpins |

**PDB search terms:**
Full Text = "capsid"
Symmetry Type = "tetrahedral"

| Type (author annotated) | Identifier Entry ID | StructureData Structure Author | PubMed ID | DOI | Structure Title |
|---|---|---|---|---|---|
| Global symmetry is icosahedral | 4RR3 | Chen, R., Lyu, K. | 25492868 | 10.1074/jbc.M114.624536 | Crystal structure of a recombinant EV71 virus particle |
| Global symmetry is icosahedral | 4YVW | Chen, R., Lyu, K. | 25833050 | 10.1128/JVI.00422-15 | crystal structure of an enterovirus 71/coxsackievirus A16 chimeric virus-like particle |

| | | | | | |
|---|---|---|---|---|---|
| Viral protease, not capsid | 2GEF | Paetzel, M., Feldman, A.R., Lee, J., Delmas, B. | 16584747 | 10.1016/j.jmb.2006.02.045 | Crystal structure of a Novel viral protease with a serine/lysine catalytic dyad mechanism |
| Polyhedrin proteins | 2WUX | Ji, X., Sutton, G., Evans, G., Axford, D., Owen, R., Stuart, D.I. | 19959989 | 10.1038/emboj.2009.352 | the crystal structure of recombinant baculovirus polyhedra |
| Polyhedrin proteins | 2WUY | Ji, X., Sutton, G., Evans, G., Axford, D., Owen, R., Stuart, D.I. | 19959989 | 10.1038/emboj.2009.352 | the crystal structure of wild-type baculovirus polyhedra |
| Polyhedrin proteins | 3JVB | Coulibaly, F., Chiu, E., Metcalf, P. | 20007786 | 10.1073/pnas.0910686106 | Crystal structure of infectious baculovirus polyhedra |
| Polyhedrin proteins | 5G3X | Bunker, R.D., Chiu, E., Metcalf, P. | 28202732 | 10.1073/pnas.1609243114 | Structure of recombinant granulovirus polyhedrin |
| Polyhedrin proteins | 6YNG | Bunker, R.D. | | | MicroED structure of granulin determined from five native nanocrystalline granulovirus occlusion |
| Polyhedrin proteins | 6S2O | Buecker, R., Mehrabi, P., Schulz, E.C., Hogan-Lamarre, P. | 32081905 | 10.1038/s41467-020-14793-0 | Granulovirus occlusion bodies by serial electron diffraction |
| Polyhedrin proteins | 5G0Z | Gati, C., Bunker, R.D., Oberthur, D., Metcalf, P., Henry, C. | 28202732 | 10.1073/pnas.1609243114 | Structure of native granulovirus polyhedrin determined using an X-ray free-electron laser |
| Polyhedrin proteins | 5MND | Oberthuer, D., Chapman, H., Doerner, K., Xavier, P.L. | 28300169 | 10.1038/srep44628 | SFX structure of Cydia pomonella granulovirus using a double flow-focusing nozzle |
| Polyhedrin proteins | 3JW6 | Coulibaly, F., Chiu, E., Metcalf, P. | 20007786 | 10.1073/pnas.0910686106 | Crystal structure of AcMNPV baculovirus polyhedra |
| Global symmetry is icosahedral | 8ELD | Subramanian, S., Bergland Drarvik, S.M., Parent, K.N. | | | Bacteriophage HRP29 Icosohedral Reconstruction |
| Non-viral protein cage | 7A4F | Tetter, S., Hilvert, D. | 34112695 | 10.1126/science.abg2822 | Aquifex aeolicus lumazine synthase-derived nucleocapsid variant NC-1 (120-mer) |
| Non-viral protein cage | 7A4G | Tetter, S., Hilvert, D. | 34112695 | 10.1126/science.abg2822 | Aquifex aeolicus lumazine synthase-derived nucleocapsid variant NC-1 (180-mer) |
| Non-viral protein cage | 7A4H | Tetter, S., Hilvert, D. | 34112695 | 10.1126/science.abg2822 | Aquifex aeolicus lumazine synthase-derived nucleocapsid variant NC-2 (180-mer) |
| Archaeal peptidase | 2CF4 | Vellieux, F.M.D., Schoehn, G., Dura, M.A., Roussel, A., Franzetti, B. | 16973604 | 10.1074/jbc.M604417200 | Pyrococcus horikoshii TET1 peptidase can assemble into a tetrahedron or a large octahedral shell |
| Non-viral protein cage | 5MQ3 | Sasaki, E., Boehringer, D., Leibundgut, M., Ban, N., Hilvert, D. | 28281548 | 10.1038/ncomms14663 | Structure of AaLS-neg |
| Polyhedrin proteins | 2OH5 | Coulibaly, F., Chiu, E., Ikeda, K., Gutmann, S., Haebel, P.W., Schulze-Briese, C., Mori, H., Metcalf, P. | 17330045 | 10.1038/nature05628 | The Crystal Structure of Infectious Cypovirus Polyhedra |
| Polyhedrin proteins | 2OH6 | Coulibaly, F., Chiu, E., Ikeda, K., Gutmann, S., Haebel, P.W., Schulze-Briese, C., Mori, H., Metcalf, P. | 17330045 | 10.1038/nature05628 | The Crystal Structure of Recombinant Cypovirus Polyhedra |
| Polyhedrin proteins | 2OH7 | Coulibaly, F., Chiu, E., Ikeda, K., Gutmann, S., Haebel, P.W., Schulze-Briese, C., Mori, H., Metcalf, P. | 17330045 | 10.1038/nature05628 | The Crystal Structure of Cypovirus Polyhedra containing the Human ZIP-kinase |

**PDB search terms:**
Full Text = "cage"
Symmetry Type = "tetrahedral"

| Type (author annotated) | Identifier Entry ID | StructureData Structure Author | PubMed ID | DOI | Structure Title |
|---|---|---|---|---|---|
| Acylphosphatase | 6KRB | Chatterjee, S., Nath, S., Sen, U. | 31866010 | 10.1016/j.bbrc.2019.12.060 | High resolution crystal structure of an Acylphosphatase protein cage |
| Designed cage | 5CY5 | Cannon, K.A., Cascio, D., Park, R., Boyken, S., King, N., Yeates, T.O. | 31840320 | 10.1002/pro.3802 | Crystal structure of the T33-51H designed self-assembling protein tetrahedron |
| Ferritin | 5XGO | Cornell, T.A., Srivastava, Y., Jauch, R., Fan, R., Orner, B.P. | 28682051 | 10.1021/acs.biochem.7b00312 | The Ferritin E-Domain: Toward Understanding Its Role in Protein Cage Assembly Through the Crystal Structure of a Maxi-/Mini-Ferritin Chimera |
| Adenosine deaminase (bacterial defense) | 8HRC | Gao, Y. | 36764292 | 10.1016/j.cell.2023.01.026 | Structure of dodecameric RdrB cage |
| Designed cage | 4NWR | McNamara, D.E., King, N.P., Bale, J.B., Sheffler, W., Baker, D., Yeates, T.O. | 24870237 | 10.1038/nature13404 | Computationally Designed Two-Component Self-Assembling Tetrahedral Cage T33-28 |
| Designed cage | 4NWN | McNamara, D.E., King, N.P., Bale, J.B., Sheffler, W., Baker, D., Yeates, T.O. | 24870237 | 10.1038/nature13404 | Computationally Designed Two-Component Self-Assembling Tetrahedral Cage T32-28 |
| Designed cage | 3M4B | Tezcan, F.A., Ni, T.W. | 20721993 | 10.1002/anie.201001487 | A Zn-mediated tetrahedral protein lattice cage |
| Designed cage | 4NWQ | McNamara, D.E., King, N.P., Bale, J.B., Sheffler, W., Baker, D., Yeates, T.O. | 24870237 | 10.1038/nature13404 | Computationally Designed Two-Component Self-Assembling Tetrahedral Cage, T33-21, Crystallized in Space Group F4132 |
| Bacterial endoprotease | 4A8C | Malet, H., Canellas, F., Sawa, J., Yan, J., Thalassinos, K., Ehrmann, M., Clausen, T., Saibil, H.R. | 22245966 | 10.1038/nsmb.2210 | Symmetrized cryo-EM reconstruction of E. coli DegQ 12-mer in complex with a binding peptide |
| Designed cage | 4NWP | McNamara, D.E., King, N.P., Bale, J.B., Sheffler, W., Baker, D., Yeates, T.O. | 24870237 | 10.1038/nature13404 | Computationally Designed Two-Component Self-Assembling Tetrahedral Cage, T33-21, Crystallized in Space Group R32 |
| Designed cage | 8UF0 | Castells-Graells, R., Meador, K., Sawaya, M.R., Yeates, T.O. | | | T33-ml23 - Designed Tetrahedral Protein Cage Using Machine Learning Algorithms |
| Designed cage | 8UMP | Castells-Graells, R., Meador, K., Sawaya, M.R., Yeates, T.O. | | | T33-ml35 - Designed Tetrahedral Protein Cage Using Machine Learning Algorithms |
| Adenosine deaminase (bacterial defense) | 8HR7 | Gao, Y. | 36764292 | 10.1016/j.cell.2023.01.026 | Structure of RdrA-RdrB complex |
| Designed cage | 4NWO | McNamara, D.E., King, N.P., Bale, J.B., Sheffler, W., Baker, D., Yeates, T.O. | 24870237 | 10.1038/nature13404 | Computationally Designed Two-Component Self-Assembling Tetrahedral Cage T33-15 |
| Dps (mini-ferritin) | 6HUI | Zeth, K., Okuda, M. | 31433627 | 10.1021/acs.inorgchem.9b00301 | The structure of Dps from Listeria innocua soaked with zinc |
| Dps (mini-ferritin) | 6HV1 | Zeth, K., Okuda, M. | 31433627 | 10.1021/acs.inorgchem.9b00301 | The apo structure of Dps from Listeria innocua before soaking experiments with Zn, Co and La |
| Dps (mini-ferritin) | 6HVQ | Zeth, K., Okuda, M. | 31433627 | 10.1021/acs.inorgchem.9b00301 | The structure of Dps from Listeria innocua soaked before soaking experiments with Zn, Co and La |
| Dps (mini-ferritin) | 6HX2 | Zeth, K., Okuda, M. | 31433627 | 10.1021/acs.inorgchem.9b00301 | The structure of Dps from Listeria innocua soaked with Cobalt |
| Bacterial endoprotease | 4A8A | Malet, H., Canellas, F., Sawa, J., Yan, J., Thalassinos, K., Ehrmann, M., Clausen, T., Saibil, H.R. | 22245966 | 10.1038/nsmb.2210 | Asymmetric cryo-EM reconstruction of E. coli DegQ 12-mer in complex with lysozyme |
| Designed cage | 4QF0 | Lai, Y.-T., Yeates, T.O. | | | Structure of a 16 nm protein cage designed by fusing symmetric oligomeric domains, quadruple |

| | | | | | |
|---|---|---|---|---|---|
| Designed cage | 6OT4 | Golub, E., Esselborn, J., Bailey, J.B., Tezcan, F.A. | 31969701 | 10.1038/s41586-019-1928-2 | Bimetallic dodecameric cage design 2 (BMC2) from cytochrome cb562 |
| Designed cage | 6OT9 | Golub, E., Esselborn, J., Bailey, J.B., Tezcan, F.A. | 31969701 | 10.1038/s41586-019-1928-2 | Bimetallic dodecameric cage design 1 (BMC1) from cytochrome cb562 |
| Designed cage | 6OT7 | Golub, E., Esselborn, J., Bailey, J.B., Tezcan, F.A. | 31969701 | 10.1038/s41586-019-1928-2 | Bimetallic dodecameric cage design 3 (BMC3) from cytochrome cb562 |
| Designed cage | 6OVH | Golub, E., Subramanian, R.H., Yan, X., Alberstein, R.G., Tezcan, F.A. | 31969701 | 10.1038/s41586-019-1928-2 | Cryo-EM structure of Bimetallic dodecameric cage design 3 (BMC3) from cytochrome cb562 |
| Bacterial endoprotease | 8F0U | Harkness, R.W., Ripstein, Z.A., Di Trani, J.M., Kay, L.E. | 37282495 | 10.1021/jacs.2c11849 | Structure of a 12mer DegP cage bound to the client protein hTRF1 |
| Acylphosphatase | 4HI2 | Nath, S., Banerjee, R., Sen, U. | | | Crystal structure of an Acylphosphatase protein cage |
| Bacterial heat shock protein | 5ZUL | Bhandari, S., Suguna, K. | 30632633 | 10.1002/prot.25657 | Small heat shock protein from Mycobacterium marinum M : Form-3 |
| Bacterial endoprotease | 3PV2 | Wrase, R., Scott, H., Hilgenfeld, R., Hansen, G. | 21670246 | 10.1073/pnas.1101084108 | Structure of Legionella fallonii DegQ (wt) |
| Bacterial endoprotease | 3PV5 | Wrase, R., Scott, H., Hilgenfeld, R., Hansen, G. | 21670246 | 10.1073/pnas.1101084108 | Structure of Legionella fallonii DegQ (N189G/P190G variant) |
| Bacterial heat shock protein | 5ZS3 | Bhandari, S., Suguna, K. | 30632633 | 10.1002/prot.25657 | Small heat shock protein from M. marinum:Form-1 |
| Designed cage | 6C9K | Gonen, S., Liu, Y., Yeates, T.O., Gonen, T. | 29507202 | 10.1073/pnas.1718825115 | Single-Particle reconstruction of DARP14 - A designed protein scaffold displaying ~17kDa DARPin proteins |
| Dps (mini-ferritin) | 1QGH | Ilari, A., Stefanini, S., Chiancone, E., Tsernoglou, D. | 10625425 | 10.1038/71236 | THE X-RAY STRUCTURE OF THE UNUSUAL DODECAMERIC FERRITIN FROM LISTERIA INNOCUA, REVEALS A NOVEL INTERSUBUNIT IRON BINDING SITE. |
| Dps (mini-ferritin) | 2C6R | Cuypers, M.G., Romao, C.V., Mitchell, E., Mcsweeney, S. | 17583727 | 10.1016/j.jmb.2006.11.032 | FE-SOAKED CRYSTAL STRUCTURE OF THE DPS92 FROM DEINOCOCCUS RADIODURANS |
| Bacterial endoprotease | 3PV3 | Wrase, R., Scott, H., Hilgenfeld, R., Hansen, G. | 21670246 | 10.1073/pnas.1101084108 | Structure of Legionella fallonii DegQ (S193A variant) |
| Ferritin | 5LS9 | Baiocco, P., Trabuco, M.C., Boffi, A. | 27942679 | 10.1039/c6nr07129e | Humanized Archaeal ferritin |
| Bacterial heat shock protein | 5ZS6 | Bhandari, S., Suguna, K. | 30632633 | 10.1002/prot.25657 | Dodecameric structure of a small Heat Shock Protein from Mycobacterium marinum M: Form-2 |
| Dps (mini-ferritin) | 6SEV | Zeth, K., Okuda, M. | 31433627 | 10.1021/acs.inorgchem.9b00301 | Structure of Dps from Listeria innocua soaked with 10 mM zinc for 120 minutes |
| Dps (mini-ferritin) | 2C2J | Cuypers, M.G., Romao, C.V., Mitchell, E., McSweeney, S. | 17583727 | 10.1016/j.jmb.2006.11.032 | Crystal Structure Of The Dps92 From Deinococcus Radiodurans |
| Non-viral protein cage | 5MQ3 | Sasaki, E., Boehringer, D., Leibundgut, M., Ban, N., Hilvert, D. | 28281548 | 10.1038/ncomms14663 | Structure of AaLS-neg |
| Designed cage | 6C9I | Gonen, S., Liu, Y., Yeates, T.O., Gonen, T. | 29507202 | 10.1073/pnas.1718825115 | Single-Particle reconstruction of DARP14 - A designed protein scaffold displaying ~17kDa DARPin proteins - Scaffold |
| Dps (mini-ferritin) | 2C41 | Ilari, A., Franceschini, S., Ceci, P., Chiancone, E. | 17018059 | 10.1111/j.1742-4658.2006.05490.x | X-ray structure of Dps from Thermosynechococcus elongatus |
| Dps (mini-ferritin) | 2CLB | Gauss, G.H., Benas, P., Wiedenheft, B., Young, M., Douglas, T., Lawrence, C.M. | 16953567 | 10.1021/bi060782u | The structure of the DPS-like protein from Sulfolobus solfataricus reveals a bacterioferritin-like di-metal binding site within a Dps- like dodecameric assembly |
| Designed cage | 4ZK7 | Liu, Y., Cascio, D., Sawaya, M.R., Bale, J., Collazo, M.J., Park, R., King, N., Baker, D., Yeates, T. | 26174163 | 10.1002/pro.2748 | Crystal structure of rescued two-component self-assembling tetrahedral cage T33-31 |
